# CRISPR Screening Uncovers a Long-Range Enhancer for *ONECUT1* in Pancreatic Differentiation and Links a Diabetes Risk Variant

**DOI:** 10.1101/2024.04.26.591412

**Authors:** Samuel Joseph Kaplan, Wilfred Wong, Jielin Yan, Julian Pulecio, Hyein S. Cho, Qianzi Li, Jiahui Zhao, Jayanti Leslie-Iyer, Jonathan Kazakov, Dylan Murphy, Renhe Luo, Kushal K. Dey, Effie Apostolou, Christina S. Leslie, Danwei Huangfu

**Affiliations:** Weill Cornell Graduate School of Medical Sciences, Weill Cornell Medical College, New York, NY, 10065, USA; Developmental Biology Program, Sloan Kettering Institute, Memorial Sloan Kettering Cancer Center, New York, NY, 10065, USA; Computational and Systems Biology Program, Sloan Kettering Institute, Memorial Sloan Kettering Cancer Center, New York, NY, 10065, USA; Louis V. Gerstner Jr. Graduate School of Biomedical Sciences, Memorial Sloan Kettering Cancer Center, New York, NY, 10065, USA; Intern, Developmental Biology Program, Sloan Kettering Institute, Memorial Sloan Kettering Cancer Center, New York, NY, 10065, USA; Meyer Cancer Center, Division of Neuro-Oncology, Department of Neurology, Sandra and Edward Meyer Cancer Center, New York-Presbyterian Hospital/Weill Cornell Medicine, New York, NY, 10065, USA

## Abstract

Functional enhancer annotation is a valuable first step for understanding tissue-specific transcriptional regulation and prioritizing disease-associated non-coding variants for investigation. However, unbiased enhancer discovery in physiologically relevant contexts remains a major challenge. To discover regulatory elements pertinent to diabetes, we conducted a CRISPR interference screen in the human pluripotent stem cell (hPSC) pancreatic differentiation system. Among the enhancers uncovered, we focused on a long-range enhancer ∼664 kb from the *ONECUT1* promoter, since coding mutations in *ONECUT1* cause pancreatic hypoplasia and neonatal diabetes. Homozygous enhancer deletion in hPSCs was associated with a near-complete loss of *ONECUT1* gene expression and compromised pancreatic differentiation. This enhancer contains a confidently fine-mapped type 2 diabetes associated variant (rs528350911) which disrupts a GATA motif. Introduction of the risk variant into hPSCs revealed substantially reduced binding of key pancreatic transcription factors (GATA4, GATA6 and FOXA2) on the edited allele, accompanied by a slight reduction of *ONECUT1* transcription, supporting a causal role for this risk variant in metabolic disease. This work expands our knowledge about transcriptional regulation in pancreatic development through the characterization of a long-range enhancer and highlights the utility of enhancer discovery in disease-relevant settings for understanding monogenic and complex disease.

## Main

A spectrum of genetic causality often influences the onset and varying severity of symptoms for a disease^1^. This concept is exemplified in metabolic conditions like diabetes, where a single nucleotide alteration in a protein coding region can cause severe neonatal diabetes mellitus (NDM) or maturity-onset diabetes of the young (MODY), while the aggregation of polygenic effects is thought to confer susceptibility to type 2 diabetes (T2D)^2,3^. However, most genetic variants have moderate effects and are in unannotated non-coding genomic regions, with the affected genes typically unknown^3,4^. Therefore, there is a pressing need to understand the functional impact of disease-associated variants uncovered by genome-wide association studies (GWAS) and manage the burgeoning set of variants of unknown significance identified through the widespread adoption of high-throughput DNA sequencing.

It is hypothesized that many genetic variants affect enhancers^5^, and alterations in their sequences can affect gene expression, contributing to both Mendelian and complex disease traits. A clear illustration of this impact is shown by the limb malformation caused by a mutation in a *SHH* enhancer ∼1 Mb from the gene promoter^6^. This represents one of a handful of clinical genetics examples where enhancer mutations have a pronounced pathological impact^7^. On the opposite end of the disease severity spectrum, functional consequences of individual sequence alterations in common complex diseases are generally unknown, but a significant proportion of disease-associated genomic variations identified through GWAS are located within putative enhancers^5^. Together, these findings have contributed to the inception of the enhanceropathy concept^7^.

However, diagnosis of suspected enhanceropathies remains difficult due to the lack of functional annotation of most enhancers. While putative enhancers can be identified based on genomic features like chromatin accessibility, complementing descriptive epigenetics through scalable functional characterization in tissue-relevant settings poses a considerable challenge. One obstacle is the variable distance enhancers can have from their target genes. In our recent interrogation of developmental enhancers, we found that 41% of the enhancer identified were located >100 kb away from the target gene promoter^8^, yet practical considerations often lead many functional enhancer screens to confine their search to regions within 100 kb of the target gene promoter^9^. Consequently, the overall prevalence of long-distance enhancer-promoter regulations remains unclear despite a growing set of long-range enhancers characterized through genetic studies^10^.

Human pluripotent stem cell (hPSC) differentiation represents a powerful system for disease modeling. The integration of genetic perturbation and genomic approaches has proven instrumental for investigating the cascade of gene and enhancer activation during development and holds promise for understanding long-term health implications. We and others have shown NDM gene knockout in hPSCs recapitulates key aspects of human genetic conditions^11^. In addition, the hPSC platform has been used to study the impact of clinically observed recessive mutations in a *PTF1A* enhancer, shedding light on its role in pancreatic agenesis and NDM^12,13^. However, many cases of NDM lack a known genetic cause even with whole exome sequencing, underscoring the importance of identifying developmental enhancers^14^. Significant progress in biochemical characterization has narrowed the search space for enhancers, but many putative enhancers have not been functionally interrogated. Furthermore, the extensive epigenetic rewiring during development further swells the number of potential enhancers, adding complexity to the characterization task^15,16^. This challenge is becoming more tractable with the use of CRISPR-Cas9 and dCas9-based CRISPR interference (CRISPRi) tools, enabling the simultaneous perturbation of genes or enhancers en masse^9^. We and others have applied CRISPR-Cas9 in hPSCs to uncover genes important for gastrulation and pancreatic development^17–22^. Recent CRISPRi screens have further identified enhancers involved in hPSC endoderm and cardiac differentiation^8,23,24^. These advances motivated us to leverage our epigenetic characterization of the hPSC pancreatic differentiation system^25^ to uncover enhancers with potential roles in diabetes.

The hPSC pancreatic differentiation system directs cells progressively through the definitive endoderm (DE), gut tube (GT), and posterior foregut (PFG, also known as PP1) stages that model embryonic development in humans^26^. Building on our recent success in using a cell identity gene as a readout for enhancer discovery in cell state transitions^8^, we utilized the developmental expression initiation of diabetes-associated gene *PDX1* at the PFG stage as a readout for the onset of pancreatic differentiation^11,17^. We then selected candidate transcription factors (TFs) likely to influence PDX1+ identity commitment and hypothesized that repressing enhancers of these TFs would affect PDX1 expression. Our subsequent screen indeed uncovered enhancers of multiple TFs that affected PDX1 activation during pancreatic specification. We focused on a distal enhancer of *ONECUT1*, located ∼664 kb from the gene promoter. This choice was motivated by the presence of a high-confidence fine-mapped T2D-associated single nucleotide polymorphism (SNP) within this enhancer^27^ and clinical genetics findings indicating that coding mutations in *ONECUT1* cause NDM^28,29^. We show that *ONECUT1* enhancer deletion in hPSCs resulted in an almost complete absence of *ONECUT1* transcript and protein, along with a reduced proportion of cells expressing PDX1 during pancreatic differentiation. Enhancer deletion had locus-wide effects within the topologically associating domain (TAD), with reduced antisense transcription and decreased H3K27ac levels. Finally, we generated hPSC lines harboring the T2D variant (rs528350911) and found the risk variant significantly impaired the binding of key pancreatic transcription factors GATA4, GATA6, and FOXA2, but had a smaller effect on *ONECUT1* transcription compared to enhancer knockout. Our study not only identified an essential enhancer of *ONECUT1* in pancreatic development but also developed and reduced to practice a framework for prioritizing disease-associated variants for investigation.

## Results

### CRISPRi enhancer screen design

To discover enhancers involved in pancreatic development and diabetes, we conducted a CRISPRi screen in a hPSC pancreatic differentiation system, using PDX1 expression as a readout. PDX1 is a key marker for the initiation of pancreatic differentiation and is indispensable for mammalian pancreatic development^30–33^. Homozygous and heterozygous mutations in *PDX1* cause NDM and maturity onset diabetes of the young (MODY), respectively^34,35^. Additionally, genomic variants in the *PDX1* locus are associated with elevated T2D risk^36^. To identify putative enhancer regions for study, we first selected 23 candidate TFs (including PDX1) based on their likely impact on PDX1+ induction and diabetes by integrating findings from murine experiments^30,31,37–55^, clinical genetics, and our previous CRISPR-Cas9 screens in hPSC pancreatic differentiation (Fig. 1B). 11 of these TFs have known roles in human metabolic disorders or NDM^29,34,56–61^. 15 TFs, including GATA6, RFX6, and ONECUT1, have also been shown to affect pancreatic differentiation and PDX1 expression in hPSC disease models or CRISPR-Cas9 coding screens for regulators of DE and PFG specification^17,18,28,62,63^.

**Fig. 1:**
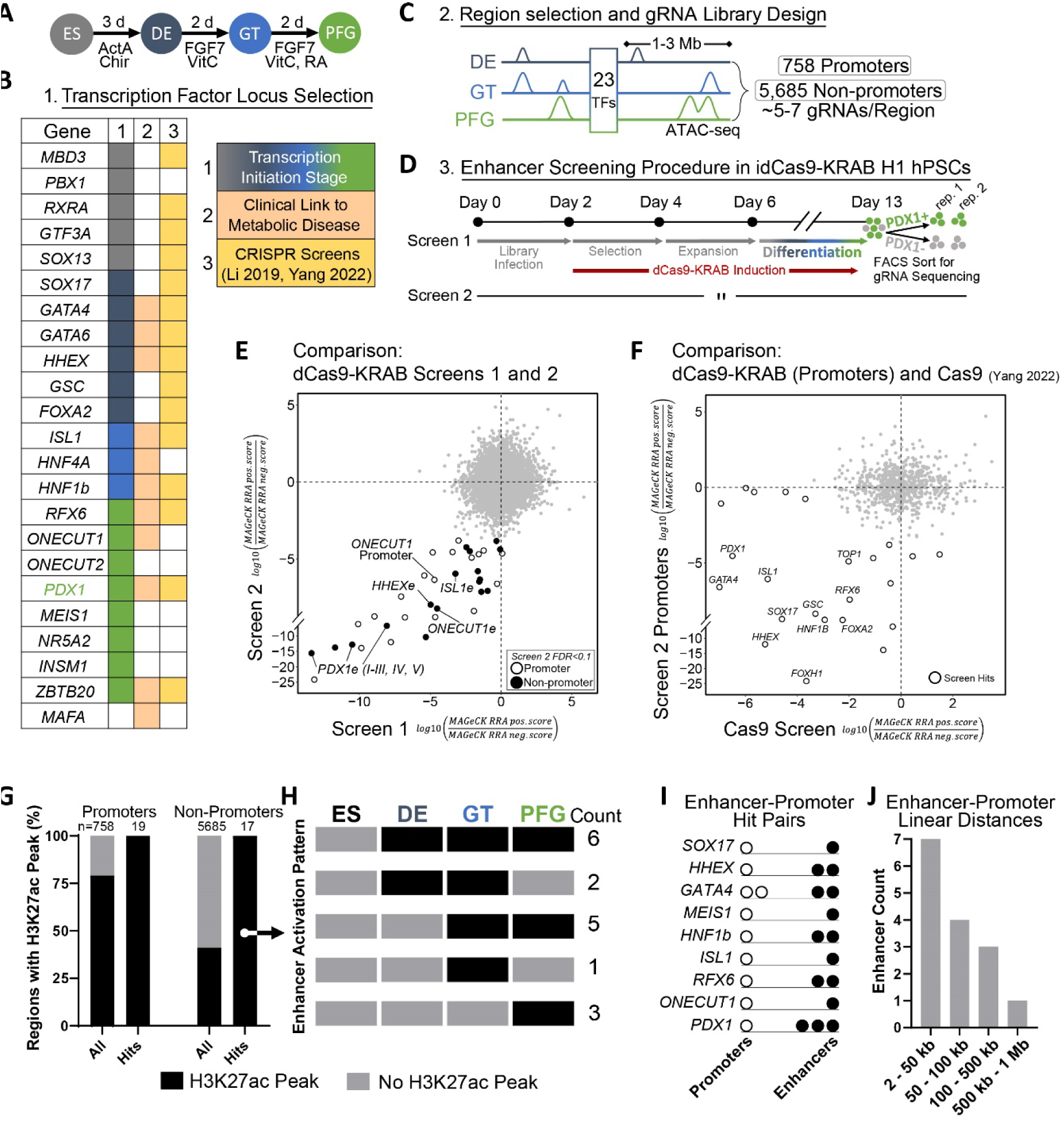
CRISPRi repression screen to discover pancreatic differentiation enhancers. A) hPSC stepwise pancreatic differentiation protocol schematic. ActA, Activin A; CHIR, CHIR99021; RA, retinoic acid; VitC, vitamin C. B) Gene selection rationale for enhancer discovery screen. C) Putative enhancer region selection and gRNA design schematic. D) Screening procedure schematic. E) Results from two screen replicates, each point represents a genomic region. Highlighted are 36 regions with a FDR < 0.1 in replicate 2, and have a decrease in PDX1 in both replicates. F) Results from screen replicate 2 compared to whole genome PDX1 expression differentiation screen. Genes showing enrichment in both screens are labeled. G) Region overlap with H3K27ac peaks at any stage during differentiation (ES, DE, GT, PFG). H) Differentiation stages at which non-promoter screen hits overlap with H3K27ac peaks, and the number of hits with each activation pattern. I) Enhancer-gene pair assignment. J) Enhancer-gene pair linear distances.

TADs have been proposed to play a key role in enhancer function^64^, so we conducted Hi-C assays on PFG stage cells and used TAD information to delineate a ∼2-6 Mb linear window around each TF for interrogation (Sup. Table 1). Within each window, we collated all accessible regions identified based on ATAC-seq peaks across the DE, GT, and PFG stages for interrogation (Fig. 1C, Sup. Table 1)^25^. These accessible regions also included 758 annotated promoters, facilitating benchmarking of the CRISPRi screen results against our previous genome-scale CRISPR-Cas9 coding screen conducted in the same differentiation context^17^.

Thus, we designed a library with 37,184 gRNAs targeting 6,443 regions (avg. length of 300 bp), 1100 safe-targeting^65^, and 463 non-targeting gRNAs^66^, and cloned the sequences into a lentiviral backbone (Sup. Table 1).

### CRISPRi screen identifies enhancers and genes necessary for human pancreatic differentiation

We conducted a pooled CRISPRi screen in our hPSC pancreatic differentiation system^17^, with two biological replicates (independent infections and differentiations) referred to as Screen 1 and 2 (Fig. 1A). For each screen, we infected inducible dCas9-KRAB H1 hPSCs^67^ with the gRNA library and induced dCas9-KRAB expression. Following differentiation, we enriched for PDX1+ and PDX1**-** cells at the PFG stage through fluorescence-activated cell sorting (FACS) (Fig. 1D, Sup. Fig. 1A). Subsequently, next-generation sequencing determined gRNA abundance within the sorted populations, and MAGeCK analysis^68^ led us to prioritize 36 top hits that caused a PDX1 decrease (Fig. 1E, Sup. Fig 1B, Sup. Table 1). Both promoter (19) and non-promoter (17) hits have greater sequence conservation compared to all regions investigated, as shown by the average conservation scores calculated from PhyloP100^69^ (Sup. Fig. 1C). Furthermore, we examined all interrogated promoter regions and found the gene hits to be largely concordant with our previous Cas9 coding screen conducted in a similar pancreatic differentiation context^17^ (Fig. 1F). These findings support the utility of CRISPRi repression for the discovery of both coding and non-coding regulators of hPSC differentiation.

We next focused on non-promoter hits and examined H3K27ac, a histone mark associated with active enhancers^70^. Non-promoter hit regions did not have H3K27ac peaks in undifferentiated hPSCs, but gained H3K27ac at various stages during pancreatic differentiation (Fig. 1G,H), indicating stage-specific enhancer activity. We further reasoned that each enhancer hit is likely to correspond to a promoter hit for the regulated gene. Indeed, almost all discovered enhancers were linked to a single promoter hit within the linear neighborhood, resulting in a total of 15 enhancer-gene pairs (Fig. 1I, Sup. Fig. 1D). This approach demonstrates the specificity of the enhancer-gene regulatory mechanism. In addition, we observed that the enhancer-promoter distances ranged from 2 to 664 kb, with three enhancers found beyond 100 kb from the transcription start site (TSS) (Fig. 1J). Thus, unbiased screening in hPSC pancreatic differentiation uncovered many previously undiscovered enhancers and expanded our understanding of gene regulation during human pancreas development.

### *ONECUT1e-664kb* deletion causes allele-specific loss of *ONECUT1* expression

The CRISPRi screen identified the *ONECUT1* promoter as well as a site ∼664 kb from the *ONECUT1* TSS, denoted as *ONECUT1e-664kb* henceforth. We observed that at the PFG stage, the *ONECUT1* promoter showed significantly increased Hi-C contact with a region encompassing *ONECUT1e-664kb* compared to the DE stage (Fig. 2A, 2B). This observation suggests a physical enhancer-promoter interaction and substantial enhancer rewiring during differentiation. We chose *ONECUT1e-664kb* for further investigation because *ONECUT1* mutations cause pancreatic hypoplasia in humans^28,29^ and *ONECUT1* is necessary for efficient hPSC differentiation to pancreatic progenitors^28^. Consistent with the clinical findings, *ONECUT1* null mice had a reduced number of endodermal cells expressing PDX1 at pancreatic budding^55^ accompanied by compromised pancreatic organogenesis^71–73^. To validate the regulatory impact of *ONECUT1e-664kb* on *ONECUT1*, we used CRISPR-Cas9 deletion as an orthogonal method to complement our dCas9-KRAB based repression approach. Using paired gRNAs flanking a ∼750 bp region, we generated two heterozygous and one homozygous clonal enhancer deletion lines (Fig. 2C). *ONECUT1* transcription is activated during the GT to PFG transition (Sup. Fig. 2A), so we differentiated the homozygous and heterozygous enhancer knockout (eKO) lines to the PFG stage and assessed PDX1 and ONECUT1 protein levels via flow cytometry (Fig. 2D). Homozygous eKO led to a near-complete depletion of ONECUT1+ cells compared to unedited cells (Fig. 2E). In addition, the percentage of cells activating PDX1 expression was significantly lower in homozygous eKO cells, showing that the loss of ONECUT1 impaired pancreatic differentiation (Fig. 2F). These findings were further confirmed through immunofluorescence staining for PDX1 and ONECUT1 (Sup. Fig. 2B). To assess the impact of enhancer knockout at a later pancreatic differentiation stage, we differentiated eKO cells to pancreatic progenitors (PP2) (Sup. Fig. 2C). Homozygous enhancer knockout cells remained ONECUT1 negative at the PP2 stage (Sup. Fig. 2F,G), and had a significant decrease in the percentage of PDX1+NKX6.1+ cells (Sup. Fig. 2D, E). This recapitulates the previously observed phenotype of *ONECUT1* gene knockout in hPSC differentiation^28^. To explore the mechanisms by which *ONECUT1* may be affecting *PDX1* expression, we examined the *PDX1* enhancers uncovered in our screen for ONECUT1 binding^17^. While all *PDX1* enhancers gained ATAC-seq and H3K27ac ChIP-seq signals during differentiation, we observed ONECUT1 ChIP-seq signal primarily at *PDX1e-2kb*, suggesting ONECUT1 may activate *PDX1* through this enhancer (Sup. Fig. 2I). Additionally, this enhancer region has binding of multiple other transcription factors such as HHEX, FOXA2, GATA4, and GATA6. This is a potential mechanism by which the cells are able to overcome loss of ONECUT1 and activate PDX1 at the PP2 stage.

**Fig. 2:**
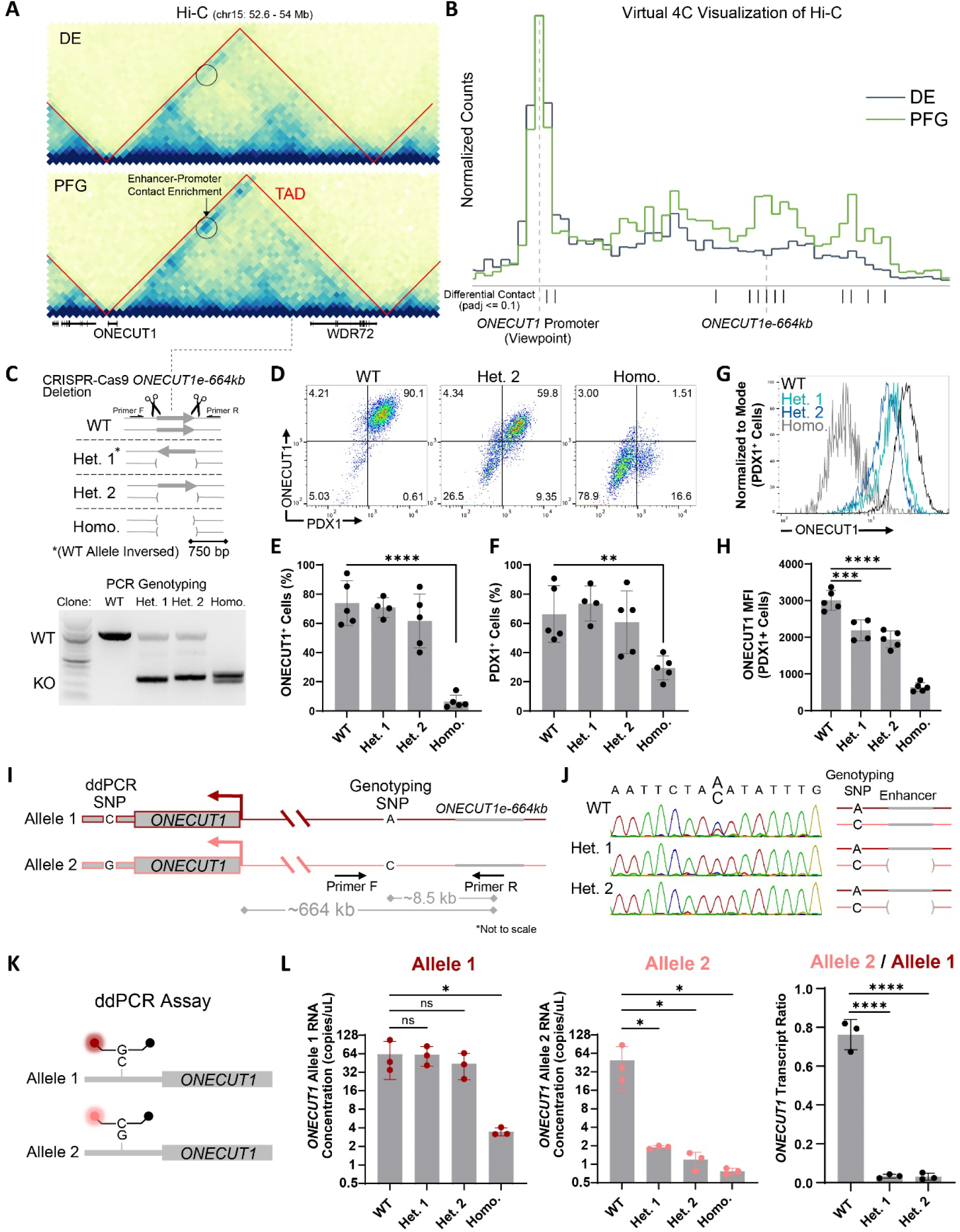
*ONECUT1e-664kb* deletion and pancreatic differentiation characterization. A) Hi-C data of definitive endoderm and posterior foregut stage differentiated hPSCs are shown around the ONECUT1 locus. Contact between ONECUT1e-664kb and ONECUT1 promoter regions is circled to highlight the increase in contact frequency between the two loci. Red lines demarcate TAD boundaries identified at 50 kb. B) 1D slices of the 2D contact map were obtained with the ONECUT1 promoter set as the bin of origin. Hi-C data was library scaled and shows the changes observed in (A). Significantly different bins were determined by DESeq2. C) hPSC enhancer deletion and genotyping PCR schematic. D) Representative flow cytometry plots of WT and enhancer knockout cells differentiated to the PFG stage. E) Percentage of cells achieving a ONECUT1+ identity. For panels E, F, and H, each dot represents one independent experiment (n = 4 or 5 independent experiments, 4 for Het. 1) and data are presented as the mean ± s.d. One-way analysis of variance (ANOVA) followed by Dunnett multiple comparisons test versus WT control. F) Percentage of cells achieving a PDX1+ identity. G) Representative histograms of ONECUT1 MFI of PDX1+ cells from flow cytometry plots of unedited and enhancer knockout cells differentiated to the PFG stage. H) Quantification and statistical comparison of ONECUT1 MFI of PDX1+ cells. I) Schematic for identifying ddPCR SNP and Genotyping SNP. J) Spectrogram of genotyping SNP amplicon sequencing and schematic showing allele that contained the deletion in heterozygous lines. K) ddPCR assay design schematic. L) Allele-specific ONECUT1 transcript quantification with ddPCR on RNA extracted from PFG stage cells. Each allele is shown in a separate plot, with the expression ratio shown in the last plot. Each symbol represents one independent differentiation (n = 3 independent experiments), with two averaged technical ddPCR replicates. Data are presented as the mean ± s.d. One-way analysis of variance (ANOVA) followed by Dunnett multiple comparisons test versus WT control.

While heterozygous lines did not exhibit a significantly lower percentage of PDX1+ or ONECUT1+ cells, there was a significant decrease in the mean fluorescence intensity signal of ONECUT1 among the PDX1+ cells in heterozygous cells compared to wild-type (WT) cells at both the PFG and PP2 stages (Fig. 2G,H, Sup. Fig. 2H). This suggested that the heterozygous cells had lower *ONECUT1* expression compared to WT cells. To directly assess the *cis* effect of heterozygous eKO on *ONECUT1* transcription, we pursued a more quantitative approach and designed an allele-specific droplet digital PCR (ddPCR) assay for the *ONECUT1* transcript.

Given the substantial linear genomic distance between *ONECUT1e-664kb* and the *ONECUT1* gene, we first called and phased all SNPs in the *ONECUT1* locus after conducting targeted long-read Nanopore sequencing with adaptive sampling (Sup. Fig. 2J). This approach allowed us to identify a SNP in the 3’UTR of *ONECUT1* and design an allele-specific ddPCR assay (Fig. 2I, K). We also identified a genotyping SNP close to *ONECUT1e-664kb* for phasing the deleted enhancer in the heterozygous lines via PCR and found both heterozygous lines had enhancer deletions on the same allele (designated as “Allele 2”) (Fig. 2J). ddPCR assays performed on RNA extracted from cells differentiated to the PFG stage showed that homozygous eKO lines lost >95% of transcription from both alleles (Fig. 2L). Heterozygous lines exhibited a specific loss of transcription from Allele 2, resulting in a significantly skewed ratio of transcripts from Allele 2 over Allele 1. This skewed ratio was not observed in control ddPCR assays conducted on genomic DNA (gDNA) (Sup. Fig. 2K). These findings demonstrate that the transcriptional impact of *ONECUT1e-664kb* deletion on *ONECUT1* expression is due to direct *in cis* regulation, rather than indirect effect through the downregulation of the ONECUT1 TF or other mechanisms. Notably, one of the heterozygous eKO lines (Het. 1) had the enhancer sequence inverted on Allele 1, and *ONECUT1* transcription from Allele 1 was not significantly affected by this inversion. Together, these results establish *ONECUT1e-664kb* as a critical *cis-*regulatory element for *ONECUT1* activation in pancreatic development.

### *ONECUT1* enhancer deletion causes transcriptional and epigenomic changes in the locus

The large impact of *ONECUT1e-664kb* deletion on *ONECUT1* transcription led us to investigate its effects on transcription in the broader genomic locus since some enhancers may serve as “hubs” that simultaneously regulate the expression of multiple genes in a locus^74^. We performed TruSeq Stranded Total (Ribodepletion) RNA-seq on WT, heterozygous, and homozygous eKO cells differentiated to the GT and PFG stages. Consistent with our ddPCR results, homozygous eKO cells had an approximately 30-fold decrease in *ONECUT1* reads at the PFG stage compared to reads in WT cells. While we did not detect any significant transcription changes of genes in the adjacent TADs, we did observe a statistically significant decrease in *WDR72* expression in homozygous *ONECUT1e-664kb* cells (Fig. 3A). Notably, *WDR72* is the only other protein-coding gene in the same TAD as *ONECUT1*. However, ddPCR assays did not show a statistically significant difference in *WDR72* transcripts, despite a noticeable trend towards a decrease (Sup. Fig. 3A), which may be attributed to the low expression of *WDR72* in the PFG stage. In summary, we conclude *ONECUT1e-664kb* primarily regulates *ONECUT1* transcription during pancreatic differentiation.

**Fig. 3:**
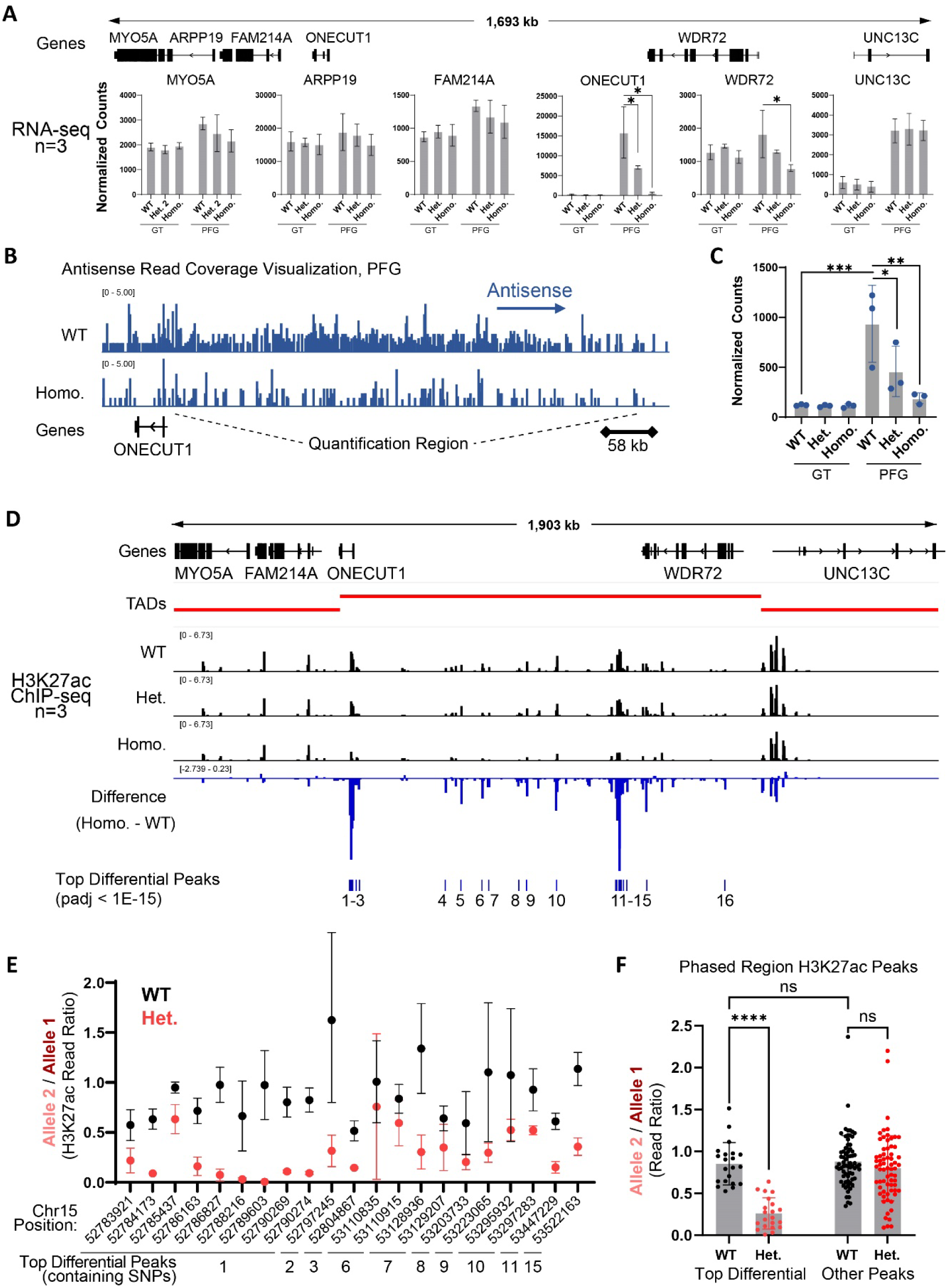
Transcriptional and epigenetic consequences of *ONECUT1e-664kb* knockout. A) RNA-seq normalized read counts for genes in ONECUT1 locus at GT and PFG stages from WT, heterozygous (het. 2), and homozygous enhancer knockout hPSCs. * indicates padj<0.005 computed by DESeq2. B) Visualization of antisense RNA relative to ONECUT1. Read coverage from strand-specific RNA-seq of PFG-stage WT and homozygous enhancer knockout cells of representative differentiation replicate. C) Quantification of (B). Each dot represents one independent experiment (n = 3 independent experiments) and data are presented as the mean ± s.d. One-way analysis of variance (ANOVA) followed by Dunnett multiple comparisons test versus WT control. D) H3K27ac ChIP-seq at PFG stage of WT, heterozygous, and homozygous enhancer knockout cells. TADs called from PFG stage HiC data. Difference calculated via MACS2 subtract. Top differential peaks between WT and homozygous enhancer knockout cells quantified with DESeq2. ONECUT1e-664kb deletion region is within peak #12. E) H3K27ac is decreased in an allele-specific manner within significantly affected H3K27ac peaks. F) Quantification of average H3K27ac read ratios at all heterozygous SNP positions in phased locus in top significantly different peaks (WT vs Homo, padj<1E-15) and other peaks. Each point represents a heterozygous SNP position, average of three replicates from independent experiments. Ordinary one-way ANOVA with Šidák’s multiple comparisons test, with a single pooled variance.

We further investigated the impact of *ONECUT1e-664kb* deletion on *ONECUT1* promoter antisense (PAS) RNA transcription. Transcription-associated PAS RNAs have been extensively documented^75,76^, and antisense RNAs can be quantified in strand-specific RNA-seq datasets^77^. Indeed, we observed an adjacent non-coding region antisense to *ONECUT1* that had lower antisense transcription in homozygous eKO cells compared to WT cells (Fig. 3B). Quantifying antisense transcripts in this region across three differentiation replicates showed that during the GT-PFG transition, *ONECUT1* transcriptional activation in WT cells was associated with a significant increase in *ONECUT1* antisense transcripts. These transcripts were significantly reduced in both heterozygous and homozygous eKO lines at the PFG stage (Fig. 3C). In contrast, sense transcripts across the same non-coding region showed no change during the GT-PFG transition and were unperturbed upon enhancer deletion (Sup. Fig. 3B). Thus, enhancer deletion has transcriptional effects beyond the decrease of the *ONECUT1* transcript, illustrating an additional transcriptional phenotype of enhancers loss.

The transcriptional changes we observed upon enhancer deletion motivated us to investigate the effect of enhancer deletion on H3K27ac: an epigenetic mark associated with active enhancers and transcription. Additionally, a previous study documented that *PTF1A* enhancer deletion caused H3K27ac loss at the *PTF1A* promoter as well as adjacent regions in the genomic locus^12^. However, given that *PTF1A* encodes a TF, it is challenging to distinguish *trans* and *cis* effects of enhancer deletion. We reasoned that analyzing allele-specific H3K27ac ChIP-seq reads could aid in examining *cis*-consequences of enhancer deletion, with a *cis-*effect manifesting as an imbalance in allele-specific reads within heterozygous eKO cells. We performed H3K27ac ChIP-seq on PFG cells differentiated from WT, heterozygous, and homozygous eKO lines. First, we compared WT and homozygous eKO cells. The decrease in H3K27ac in the *ONECUT1* TAD was the most pronounced of all regions on chromosome 15 (Sup. Fig. 3C), strongly suggesting *cis* effects. Among the 16 regions with the most significant H3K27ac reduction (adjusted p-value < 1E-15) (Fig. 3D), 10 contained heterozygous SNPs. Since these SNPs were phased with long-read sequencing (Sup. Fig. 2J), we could use the SNPs to quantify changes occurring *in cis* with enhancer deletion by calculating allele-specific H3K27ac ChIP-seq read ratios in both WT and heterozygous eKO cells (Sup. Fig. 3D). As controls, we used regions with heterozygous SNPs that did not meet the significance threshold of the differential analysis. For all top differential peaks, H3K27ac was decreased *in cis* to the allele with enhancer deletion in heterozygous eKO cells, with some positions losing nearly all H3K27ac (Fig. 3E), whereas control regions showed a near 1:1 allele read ratio average in both WT and heterozygous eKO cells (Fig. 3F). This demonstrates *ONECUT1e-664kb* affects H3K27ac at multiple genomic positions within the locus, supporting a model of direct crosstalk among the putative regulatory elements. Thus, the reduction of H3K27ac signal and PAS transcripts *in cis* with enhancer deletion illustrates the impact of enhancer dysregulation on genome function beyond gene transcription.

### T2D-associated SNP decreases pancreatic TF recruitment to *ONECUT1e-664kb*

To explore the significance of *ONECUT1e-664kb* in a clinical genetics context, we examined SNP metabolic disease-association statistics in the *ONECUT1* locus using data from a recent T2D GWAS meta-analysis^27^ and other studies^78,79^. We observed one distinct T2D disease association signal around the *ONECUT1* gene with a largely unresolved 99% credible set spanning ∼100kb genomic region (hg38, chr15:52777944-52873484). We identified a second independent T2D disease-causal signal ∼664 kb away from *ONECUT1* and close to *WDR72*, that is driven by a single variant, rs528350911 (G/C, reverse strand, minor allele frequency = 0.0068)^27^ (Fig 4A). This variant is directly within *ONECUT1e-664kb* and is highly confidently fine-mapped with the posterior probability of association (PPA) close to 1 (0.994) after linkage disequilibrium (LD) correction that is part of SuSiE fine-mapping^27,80^. To predict if rs528350911 would affect enhancer activation in pancreatic development, we trained a ChromBPNet^81,82^ model on PFG-stage ATAC-seq data^25^. Then, we computed per nucleotide DeepLift^83^ attribution scores across the region for both the reference and alternate sequences of rs528350911. The decreased attribution for an entire GATA TF motif suggests that the G in the GATA motif is crucial for the model’s genomic accessibility prediction, and the alternate sequence associated with T2D risk leads to an attribution loss at the site (Fig. 4B).

**Fig. 4:**
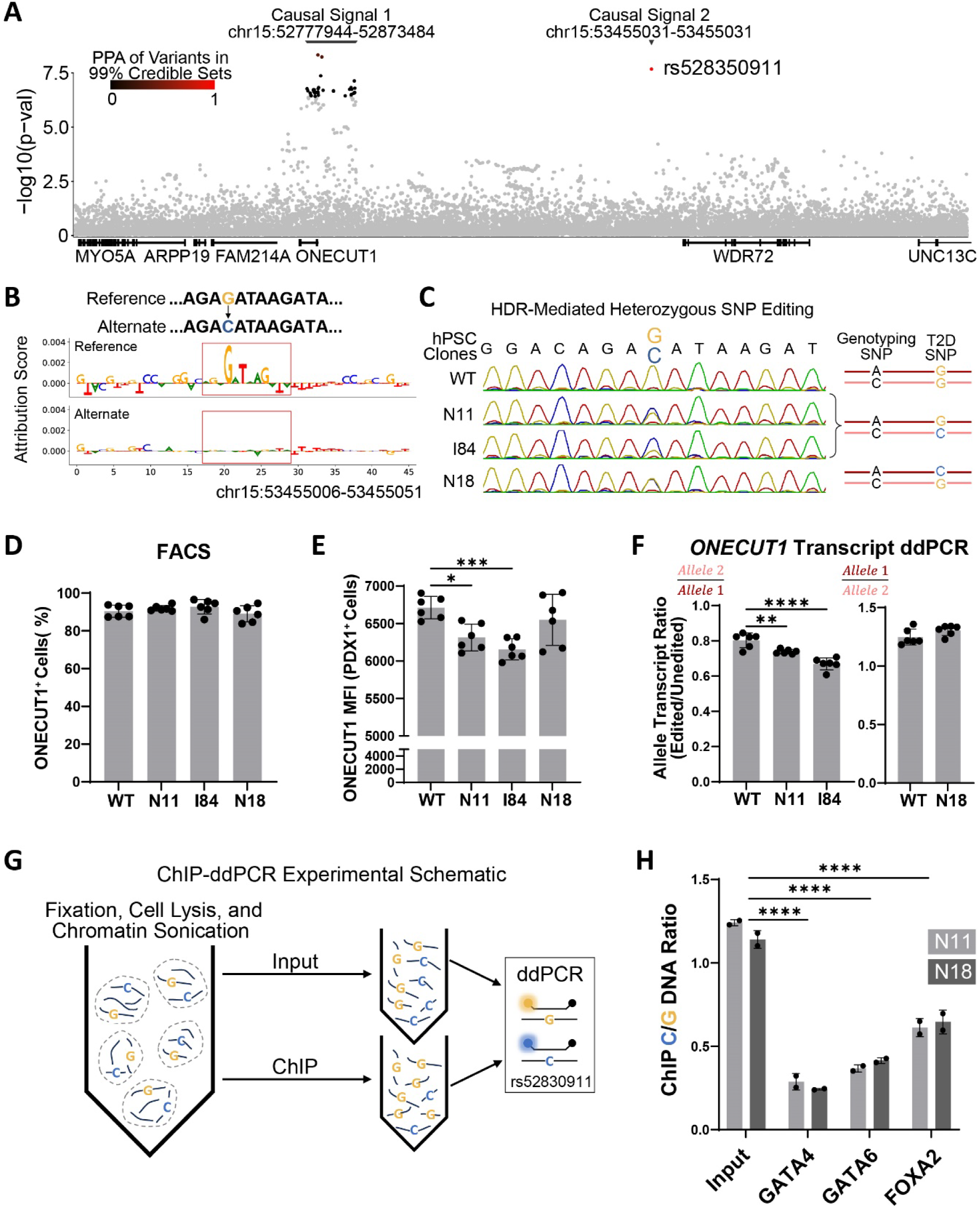
T2D-associated SNP hPSC modeling and pancreatic differentiation characterization. A) LocusZoom plot of variants from Mahajan et al (2018b) T2D GWAS meta-analysis unadjusted for BMI. Variants within credible sets are colored by the PPA score. B) Decreased DeepLift attribution score at GATA4 motif upon introduction of variant. Red box represents a GATA4/6 binding motif hit by FIMO (p < 0.001). C) Spectrogram of disease-associated SNP amplicon sequencing and schematic demonstrating allele that contains the introduced SNP in hPSCs. D) Percentage of ONECUT1+ cells at PFG stage. For panels D, E, and F, each symbol represents one independent experiment (n = 6 independent experiments) and data are presented as the mean ± s.d. One-way analysis of variance (ANOVA) followed by Dunnett multiple comparisons test versus WT control. E) Statistical comparison of ONECUT1 MFI of PDX1+ cells. F) ONECUT1 transcript ratio of edited allele divided by unedited allele. For each independent experiment shown, two averaged technical ddPCR replicates. G) ChIP-ddPCR schematic. H) ChIPed DNA ratio analysis. Each symbol represents one independent differentiation and ChIP experiment, two averaged technical ddPCR replicates. Two-way analysis of variance (ANOVA) followed by Tukey’s multiple comparisons test versus WT control.

GATA factors such as GATA4 and GATA6 are known for their important role in pancreatic differentiation^62^ and are linked to neonatal diabetes^84,85^. At the PFG stage, the key pancreatic TFs GATA4, GATA6, and FOXA2, bind to the *ONECUT1e-664kb* enhancer region (Sup. Fig. 4A)^25,62^. This motivated us to investigate the molecular impact of rs528350911. We engineered three heterozygous hPSC lines with the risk variant: two carried the variant on Allele 2, and one on Allele 1 (Fig. 4C). While all lines differentiated to the PFG stage had a similar percentage of ONECUT1 and PDX1 expressing cells (Fig. 4D, Sup. Fig. 4B) and did not affect transcription of WDR72 (Sup. Fig. 4C), we observed a significant reduction in ONECUT1 MFI for two of the three lines (Fig. 4E). Allele-specific ddPCR quantification of *ONECUT1* transcription (Fig. 2K) confirmed that the same two lines had a small but significant allele-specific reduction in *ONECUT1* transcription (Fig. 4F).

We further investigated the impact of rs528350911 on TF binding at *ONECUT1-e664kb*. To accomplish this, we developed an allele-specific ddPCR assay targeting the reference and alternate (T2D associated) nucleotides (Fig. 4G) to quantify TF binding following chromatin immuno-precipitation (ChIP) in an allele-specific manner. By utilizing the heterozygous lines, this allele-specific assay supports an internally controlled experimental system that mitigates technical variabilities arising from *in vitro* differentiation, ChIP, and PCR. We selected two hPSC lines that contained the risk variant on different alleles; one line had a significant impact on *ONECUT1* expression, while the other did not show a significant effect as mentioned above. After differentiating the lines to the PFG stage, we performed ChIP for GATA4, GATA6, and FOXA2. The precipitated DNA was then used as the template in the ddPCR assay, revealing a ∼4-fold reduction in GATA4 binding on the allele containing the disease-associated SNP in both lines (Fig. 4H). We also observed significantly decreased GATA6 and FOXA2 binding. Thus, despite a marginal effect on *ONECUT1* transcription, the disease-associated SNP substantially reduced TF binding to *ONECUT1e-664kb*.

## Discussion

Pancreatic developmental enhancers have been found in mice by integrating pre-existing knowledge of important pancreatic genes with annotations such as evolutionary conservation and DNA accessibility. Using this approach, several murine *PDX1* enhancers were identified, and knockout mice had a spectrum of pancreatic phenotypes depending on the specific region that was removed^86,87^. A similar approach also identified a putative *GATA4* enhancer, which was subsequently verified by a transgenic reporter assay in mouse endoderm and endoderm-derived tissues including the pancreas and duodenum^88^. These studies set the stage for integration of biochemical epigenetic marks into the enhancer discovery framework, which led to the discovery of a *PTF1a* enhanceropathy responsible for pancreatic agenesis and neonatal diabetes in humans^12,13^. The studies also highlight the value of developmental enhancer discovery for elucidating mechanisms of organogenesis and disease diagnosis. Building upon these findings, our study expanded the scope of enhancer discovery by targeting developmentally accessible chromatin regions for large-scale functional interrogation using dCas9-KRAB. Our study not only identified the human *PDX1* and *GATA4* enhancers corresponding to the aforementioned mouse enhancers (Sup. Table 1), but also uncovered additional enhancers that had not been previously described. These newly discovered enhancers can serve as a diagnostic resource not only for the large number of monogenic diabetes patients with no known genetic cause^14^, but also for comprehending variants identified through population-level association studies.

From all the discovered enhancers, this work contributes *ONECUT1e-664kb* to a select group of well-documented long-range regulatory sequences that activate transcription over genomic intervals exceeding 500 kb^89–97^. These enhancers are valuable models for further investigation into long-range transcriptional regulation and its potential implications in human disease. We speculate that gene regulation by long-range enhancers is more common than appreciated, and our findings underscore the value of unbiased discovery. From a technical standpoint, the substantial genomic distance between *ONECUT1e-664kb* and its target gene presented a challenge for definitively demonstrating the *cis* impact of enhancer deletion. In addition to conducting Hi-C to support enhancer-promoter interaction, we phased the variants in the *ONECUT1* locus and employed high-sensitivity variant-specific ddPCR assays in heterozygous lines. As a result, we uncovered two mechanisms beyond transcriptional control by which regulatory regions could influence disease traits: loss of promoter antisense transcription and H3K27ac in a genomic locus. We also demonstrated a near-complete, allele-specific loss of *ONECUT1* transcription resulting from the deletion of *ONECUT1e-664kb*. Considering results from *ONECUT1* knockout studies in mice^55^ and hPSC pancreatic differentiation^28^, we infer ONECUT1 protein loss to be the driver of the pancreatic differentiation phenotype in our study. Our findings are also in concordance with the reduced number of PDX1 expressing cells during pancreatic budding in *ONECUT1* null mice that contributes to pancreatic hypoplasia^55^. We predict that loss of *ONECUT1e-664kb* in an individual would phenocopy the severe pancreatic hypoplasia observed in cases of *ONECUT1* protein-coding mutations in clinical settings^28,29^.

The discovery of *ONECUT1e-664kb* also helped us prioritize a disease risk variant for functional interrogation. In previous GWAS, the T2D-associated variant rs528350911 is assigned to the closest gene – *WDR72* ^27^. Considering the strong effect of *ONECUT1e-664kb* deletion on *ONECUT1* expression and the known association of *ONECUT1* with diabetes, it is likely this variant confers disease susceptibility through its effect on ONECUT1 expression. Our study demonstrates how the integration of functional screening, precise genome editing, deep learning models, and genomic conformation information can improve the functional pairing of variants with genes. The introduction of the disease variant into hPSCs did not have a strong effect on *ONECUT1* transcription, but its robust effect on TF binding to *ONECUT1e-664kb* suggests that assays other than gene expression could be valuable for characterizing the molecular consequences of variants in endogenous loci. We speculate rs528350911 may confer a latent disease vulnerability, which could be exposed by factors not fully modeled in our current differentiation system such as age and environment^98,99^. Given the considerable distance between *ONECUT1e-664kb* and the *ONECUT1* gene, as well as the limited impact of the rs528350911 risk variant on *ONECUT1* transcription, it is unlikely this variant would have been prioritized for investigation in a high-throughput variant modeling screen that relies on prime or base editing technologies. Assays with higher sensitivity are likely necessary to detect smaller magnitude effects of point variants like rs528350911 compared to the more pronounced effects of enhancer repression with dCas9-KRAB^100^. Our work demonstrates that integrating *in vitro* functional enhancer discovery with population-level genomics provides a rational approach to efficiently prioritize variants for deeper investigation. This framework holds promise for understanding the exponentially growing set of disease-associated variants discovered through population-wide GWAS and whole genome sequencing efforts.

## Limitations of the Study

1. We were not able to identify the promoter hits for two of the enhancers, presumably due to variations in gRNA efficiency and screen cutoff stringency. The investigation was also limited to the embryonic pancreatic differentiation system. This system may not be optimal for investigating disease-associated variants for complex diseases that develop over a person’s adult lifetime such as Type II Diabetes. Additionally, genes can have different functions in different tissues, and it is possible there are additional non-developmental^72,73^ or non-pancreatic^101^ mechanisms through which ONECUT1 variants could contribute to metabolic disorders.
2. There are long non-coding RNAs (lncRNAs) in the intergenic region between *ONECUT1* and *WDR72* with the annotations differing substantially between transcriptome assemblies. In some databases, rs528350911 has been linked to one of these lncRNAs, *LINC02490*. We hypothesize this is the case because it is the nearest annotated element centromeric to rs528350911, even though it is ∼330kb away. However, very low expression makes it challenging to quantify the impact of enhancer knockout on any individual lncRNA from our RNA-seq data. We chose to evaluate them in aggregate in the PAS analysis, however we recognize it is possible individual lncRNAs may have molecular consequences.
3. While we have generated molecular evidence for the role of rs528350911 in pancreatic differentiation, it is still plausible the SNP may not directly cause metabolic disorders and further studies are needed.

## Acknowledgments

We thank Dr. Thomas Vierbuchen for valuable advice. We thank Hanuman Kale and Emily DeBitetto for assisting with experiments not included in the manuscript and acknowledge the assistance from the following Memorial Sloan Kettering Cancer Center (MSKCC) Cores: Integrated Genomics Operation, Flow Cytometry, Gene Editing & Screening, Molecular Cytogenetics, and Stem Cell Research. We thank Dr. Ralph Garippa and Sanjoy Mehta for assistance with CRISPR library generation and sequencing, Cassidy Cobbs for providing advice regarding next-generation sequencing experiments, Dr. Stephanie Chrysanthou for conducting nanopore sequencing with adaptive sampling, and Dr. Andrea Farina for designing and executing the ddPCR assays.

## Author Contributions

Conceptualization: SJK, DH

Methodology: SJK, JY, WW, JP, HSC, RL, DM, KD, EA, CSL, DH

Investigation: SJK, JY, WW, JP, HSC, JLI, JK, JZ, QL, DM, RL

Visualization: SJK, WW, QL

Funding acquisition: DH

Project administration: DH

Supervision: SJK, DH

Writing (original draft): SJK, DH

Writing (review & editing): all authors

## STAR★Methods

### KEY RESOURCES TABLE

#### RESOURCE AVAILABILITY

- Lead contact

- Materials availability

- Data and code availability

### EXPERIMENTAL MODEL AND STUDY PARTICIPANT DETAILS

- Primary Cells

### METHOD DETAILS

- hPSC culture

- hPSC directed pancreatic differentiation

- gRNA library assembly

- Lentiviral gRNA libraries

- CRISPR dCas9-KRAB screens

- gRNA sequencing and screen analysis

- Generation of *ONECUT1e-664kb* clonal knockout hPSC lines

- Generation of clonal hPSC lines with disease-associated variant

- *ONECUT1* locus SNP phasing

- *ONECUT1e-664kb* heterozygous knockout line phasing

- *ONECUT1e-664kb* heterozygous rs528350911 knock-in line phasing

- Immunofluorescence staining

- Flow cytometry analysis

- RNA-seq and analysis

- Droplet digital PCR (ddPCR) and analysis

- Hi-C and analysis

- ChIP and ChIP-seq assays

- H3K27ac ChIP-seq analysis

- chromBPNet variant modeling

## QUANTIFICATION AND STATISTICAL ANALYSIS

KEY RESOURCES TABLE.

| REAGENT or RESOURCE | SOURCE | IDENTIFIER |
| --- | --- | --- |
| Antibodies |  |  |
| SOX17 | BD BIOSCIENCES | Cat#561591; RRID: AB_10717121 |
| CXCR4 | R&D Systems | Cat#FAB170A; RRID: AB_357073 |
| PDX1 | R&D Systems | Cat#AF2419; RRID: AB_355257 |
| ONECUT1 | Santa Cruz Biotechnology | Cat#sc-376308; RRID: AB_10988385 |
| NKX6-1 | DHSB | Cat#F55A12; RRID: AB_532379 |
| Alexa Fluor 488 | ThermoFisher Scientific | Cat#A-11055; RRID: AB_2534102 |
| Alexa Fluor 647 | ThermoFisher Scientific | Cat#A-31571; RRID: AB_162542 |
| LIVE/DEAD™ Fixable Violet Dead Cell Stain Kit | ThermoFisher Scientific | Cat#L34964 |
| FOXA2 | Cell Signaling Technology | Cat#8186; RRID: AB_10891055 |
| GATA4 | Invitrogen | Cat#MA5-15532; RRID: AB_10989032 |
| GATA6 | Cell Signaling Technology | Cat#5851; RRID: AB_10989032 |
| H3K27ac | EMD Millipore | Cat#MABE647; RRID: AB_2893037 |
| Bacterial and virus strains |  |  |
| NEB 5-alpha Competent E. coli | NEB | Cat#C2987 |
| Chemicals, peptides, and recombinant proteins |  |  |
| Complete Essential 8 (E8) medium | Thermo Fisher Scientific | A1517001 |
| Vitronectin (VTN-N) | Thermo Fisher Scientific | A14700 |
| EDTA | KD Medical | RGE-3130 |
| TrypLE Select 1X | Thermo Fisher Scientific | 12563-029 |
| ROCK inhibitor Y-27632 2HCl | Selleck Chemicals | S1049 |
| Doxycycline | Sigma | D9891 |
| Lipofectamine RNAiMAX Transfection Reagent | Thermo Fisher Scientific | 13778150 |
| Opti-MEM | Thermo Fisher Scientific | 31985070 |
| MCDB 131 | Thermo Fisher Scientific | 10372019 |
| Activin A | PeproTech | 120-14E |
| CHIR 99021 | Tocris | 442310 |
| L-Ascorbic acid (vitamin C) | Sigma-Aldrich | A4544 |
| FGF7 (KGF) | PeptoTech | 100-19 |
| SANT-1 | Sigma | S4572 |
| Retinoic acid | Sigma | R2625 |
| LDN-193189 | Reprocell | 04-0074 |
| TPB | EMD Millipore | 565740 |
| ITS-X (100X) | Thermo Fisher Scientific | 51500-056 |
| Dynabeads Protein G | Thermo Fisher Scientific | 10004D |
| Critical commercial assays |  |  |
| Herculase II Fusion DNA Polymerase | Agilent Technologies | 600679 |
| LongAmp Taq DNA Polymerase | NEB | M0323 |
| eBioscience™ Foxp3 / Transcription Factor Staining Buffer Set | Thermo Fisher Scientific | 00-5523-00 |
| Arima-Hi-C kit | Arima | A510008 |
| Monarch® HMW DNA Extraction Kit | NEB | T3050 |
| TruSeq Stranded Total RNA | Illumina | 20020597 |
| Zero Blunt TOPO PCR Cloning Kit | Thermo Fisher Scientific | 450245 |
| Deposited data |  |  |
| Raw and processed data | This paper | GSE267330 |
| Raw and processed data | Lee et al., 2019 | GSE114102 |
| Raw and processed data | Yang et. al 2022 | GSE181480 |
| Raw and Processed Data | Luo et al., 2023 | 4DNFIKCEWMIA, 4DNFI7RGXYFY, 4DNFIRLG5UWL |
| Experimental models: Cell lines |  |  |
| H1 dCas9-KRAB line | Pulecio et al., 2023 | Derived from the parental line (NIHhESC-10-0043); RRID: CVCL_WS14 |
| H1 iCas9 line | Shi et al., 2017 | Derived from the parental line (NIHhESC-10-0043); RRID: CVCL_WS14 |
| Oligonucleotides |  |  |
| See Table S1 Sheet 3 for screen gRNA spacer sequences, Table S3 for all other sequences | This paper | N/A |
| Software and algorithms |  |  |
| PRISM v10 | GraphPad Software | https://www.graphpad.com/; RRID: SCR_002798 |
| Flowjo-v10 | FlowJo LLC | https://www.flowjo.com/; RRID: SCR_008520 |
| R | 4.4.0 | https://cran.r-project.org/; RRID: SCR_003005 |
| Genrich | 0.6.1 | https://github.com/jsh58/Genrich; RRID: SCR_025320 |
| CHOPCHOPv3 |  | https://chopchop.cbu.uib.no/; RRID: SCR_015723 |
| MAGECK | 0.5.9.5 | https://sourceforge.net/p/mageck/wiki/Home/; RRID: SCR_025016 |
| Minimap2 | 2.26 | https://github.com/lh3/minimap2; RRID: SCR_018550 |
| samtools | 1.2 | http://htslib.org/; RRID: SCR_002105 |
| WhatsHap | 2.2 | https://whatschap.readthedocs.io/; RRID: SCR_025319 |
| longshot | 0.4.5 | https://github.com/pjedge/longshot; RRID: SCR_025318 |
| STAR | 2.7.11b | https://github.com/alexdobin/STAR; RRID: SCR_004463 |
| Subread featureCounts | 2.0.6 | https://subread.sourceforge.net/; RRID: SCR_009803 |
| QuantaSoft Analysis Pro | 1.0.596 | https://www.bio-rad.com; RRID: SCR_025321 |
| HiC-Pro | 3.1.0 | https://github.com/nservant/HiC-Pro; RRID: SCR_017643 |
| DESeq2 | 1.40.1 | https://bioconductor.org/packages/release/bioc/html/DESeq2.html; RRID: SCR_015687 |
| HiTC | 1.34.0 | http://www.bioconductor.org/packages//2.10/bioc/html/HiTC.html; RRID: SCR_013175 |
| HiCDCPlus | 1.5.2 | https://www.bioconductor.org/packages/release/bioc/html/HiCDCPlus.html; RRID: SCR_025317 |
| Bowtie2 | 2.5.1 | http://bowtie-bio.sourceforge.net/bowtie2/index.shtml; RRID: SCR_016368 |
| MACS2 | 2.2.9.1 | https://github.com/macs3-project/MACS; RRID: SCR_013291 |
| IDR | 2.0.3 | https://github.com/nboley/idr; RRID: SCR_017237 |
| IGV | 2.17.4 | http://www.broadinstitute.org/igv/; RRID: SCR_011793 |
| GATK | 4.0.1.2 | https://software.broadinstitute.org/gatk/; RRID: SCR_001876 |
| ChromBPNet | 0.1.7 | https://github.com/kundajelab/chrombpnet; RRID: SCR_024806 |
| MEME Suite (FIMO) | 5.5.5 | http://meme-suite.org/; RRID: SCR_001783 |
| bedtools | 2.31.1 | https://github.com/arq5x/bedtools2; RRID: SCR_006646 |
| Other |  |  |

## RESOURCE AVAILABILITY

### Lead Contact

Further information and requests for resources and reagents should be directed to Danwei Huangfu.

### Materials availability

All the hPSC cell lines and plasmids generated in this study are available upon reasonable request.

### Data and code availability

- All the sequencing data generated in this study are available at GEO under accession code GSE267330. The Hi-C data from the PFG cells are also available in the 4DN Nucleome Data Portal (https://data.4dnucleome.org) under accession number 4DNFIQBPRZZD. Previously published data that were re-analyzed here are available at GEO under codes GSE114102 (ATAC-seq data used for screen design, and ATAC-seq and H3K27ac ChIP-seq visualization in Sup. Fig. 1E) and GATA4, GATA6, and FOXA2 ChIP-seq) and GSE181480 (HHEX and ONECUT1 ChIP-seq), and the 4DN Nucleome Data Portal, accession numbers: 4DNFIKCEWMIA, 4DNFI7RGXYFY, 4DNFIRLG5UWL. All genomic coordinates reported in manuscript are hg38 unless otherwise indicated.
- This paper does not report original code.
- Any additional information required to reanalyze the data reported in this work paper is available from the lead contact upon request.

## EXPERIMENTAL MODEL AND STUDY PARTICIPANT DETAILS

### Primary cells

The parental hESC line H1 (NIHhESC-10-0043) is available from WiCell under a material transfer agreement.

## METHOD DETAILS

### hPSC culture

Experiments were performed with H1 human embryonic stem cells (NIHhESC-10-0043), which were regularly confirmed to be mycoplasma-free by the Memorial Sloan Kettering Cancer Center (MSKCC) Antibody & Bioresource Core Facility. All experiments were conducted per NIH guidelines and approved by the Tri-SCI Embryonic Stem Cell Research Oversight (ESCRO) Committee. hPSCs were maintained in Essential 8 (E8) medium (Thermo Fisher Scientific, A1517001) on vitronectin (Thermo Fisher Scientific, A14700) pre-coated plates at 37°C with 5% CO_2_. The Rho-associated protein kinase (ROCK) inhibitor Y-276325 (5 µM; Selleck Chemicals, S1049) was added to the E8 medium immediately after passaging or thawing of hPSCs for 24 hours.

### hPSC directed pancreatic differentiation

hPSCs were maintained in E8 medium for 2 days to reach ∼80% confluence. Cells were washed with PBS and differentiated to DE, GT, PFG, and PP2 stages following previously described protocols^17,102^. hPSCs were rinsed with PBS and first differentiated into DE using S1/2 medium supplemented with 100 ng/ml Activin A (Bon Opus Biosciences) for 3 days and CHIR99021 (Stemgent, 04-0004-10) for 2 days (first day, 5 µM; second day, 0.5 µM). DE cells were rinsed with PBS and then exposed to S1/2 medium supplemented with 50 ng/ml KGF (FGF7) (PeproTech, 100-19) and 0.25 mM vitamin C (VitC) (Sigma-Aldrich, A4544) for 2 days to reach GT stage. GT cells were then switched to S3/4 medium supplemented with 50 ng/ml FGF7, 0.25 mM VitC and 1 µM retinoic acid (RA) (Sigma-Aldrich, R2625) for 2 days to reach PFG stage. PFG cells were then switched to S3/4 medium supplemented with 2 ng/ml FGF7, 0.25 mM VitC, 0.1 µM RA, 200 nM LDN, 0.25 µM SANT-1, 100 nM TPB and 1:200 ITS-X for 4 days to reach PP2 stage.

### gRNA library assembly

23 transcription factors expressed in pancreatic differentiation were selected based on known clinical roles in metabolic disease and demonstrated pancreatic functional requirement from CRISPR screens^29,56–59^. Around each transcription factor, loci within hg19 (lifted over in supplemental tables to hg38) were defined to include at least the TAD containing the gene, with TADs called on PFG-stage Hi-C data as previously done^67^. To select putative enhancers for interrogation in these loci, Genrich (0.6.1)^103^ was used to call peaks at an FDR-adjusted p-value of 0.01 on previously published DE, GT, and PFG stage ATAC-seq data (Sup. Table 1)^25^. For each region in the loci, and some additional regions, CHOPCHOPv3^104^ was used to design gRNAs. For regions <750 bp, the top 5 ranked gRNAs were taken, and for regions >750 bp, 1 gRNA/150bp of sequence was selected for interrogation. Finally, 1100 safe harbor^65^ and 463 non-targeting gRNAs^66^ were included in the library for a total of 38,747 gRNAs. Oligos were synthesized, amplified, and restriction cloned into the lentiGuide-puro^105^ (Addgene: 52963; RRID: Addgene_52963) backbone by the MSKCC Gene Editing & Screening Core Facility.

Cloned plasmid libraries were PCR amplified to incorporate adapters for NGS. Samples were purified and sequenced using Illumina HiSeq 2500 platform. FASTQ files were clipped by position and reads were mapped back to the reference library file to quantify abundance of reads per gRNA. The overall representation of the libraries was charted over a one-log fold change to evaluate representation in the final library.

### Lentiviral gRNA libraries

gRNA lentiviral library generation was performed as previously described^17^. In brief, a total of 9.45 μg of the library plasmid combined with 6.75 μg lentiviral packaging vector psPAX2 and 1.36 μg vesicular stomatitis virus G (VSV-G) envelope expressing plasmid pMD2.G (Addgene plasmids 12260 and 12259) were transfected with the JetPRIME (VMR; 89137972) reagent into 10E6 293T cells in a 10cm plate to produce the lentiviral particles. Fresh medium was changed 24h after transfection and viral supernatant was collected at 48 and 72h after transfection, spun at 1000 rpm for 5 minutes, 0.45um filtered, and stored at −80°C.

### CRISPR dCas9-KRAB screens

Two independent screens were performed following the same procedure. H1 hPSC (i)dCas9-KRAB cells were used^67^. The lentiviral library was transduced into 12E6 hPSCs distributed in three 10 cm plates following the conditions (MOI: 4, protamine sulfate: 6 μg/ml, Ri: 10 μM). After one day of recovery, doxycycline (DOX, 2 μg/ml) was added, and cells were kept under puromycin selection for 2 days. After two days, 30E6 cells were seeded into 6-well VTN-coated plates (5E5 per well) and grown for 48 hours in E8 medium. After washing with Phosphate Buffered Saline (PBS) w/o Ca2+ & Mg2+, cells were differentiated, with DOX added throughout the seven-day differentiation process. At the PFG stage, cells were dissociated with TrypLE, stained with LIVE/DEAD reagent, fixed, stained with the PDX1 antibody, and stained with a fluorescent secondary antibody. They were then sorted by the MSKCC flow cytometry facility based on 2ndary antibody fluorescence using FACSAria sorters (BD Biosciences).

Positive and negative cells were sorted aiming for a minimum 1000X representation (number of cells sorted*MOI)/(unique gRNAs in library) per condition, with two independent sorts serving as technical replicates for each biological replicate. For biological replicate 1, approximately 10E6 PDX1- and 10E6 PDX1+ cells were sorted. For replicate 2, approximately 20E6 PDX1- and 20E6 PDX1+ cells were sorted. Cells were pelleted and kept at -80°C for downstream gRNA sequencing.

### gRNA sequencing and screen analysis

gRNA enrichment sequencing was performed by the MSKCC Gene Editing & Screening Core Facility as previously described, but with modifications to enable DNA extraction from fixed cells^17^. Cells were decrosslinked and treated with proteinase K (0.1 mg/ml) overnight followed by genomic DNA extraction with QIAGEN Blood & Cell Culture DNA Maxi Kit (QIAGEN; 13362) and quantified by Qubit (Thermo-Scientific) following the manufacturer’s guidelines. A quantity of gDNA sufficient for 1000X representation of gRNAs was amplified with oligos containing the Illumina adapters and multiplexing barcodes by PCR. Amplicons were quantified by Qubit and Bioanalyzer (Agilent) and sequenced on the Illumina HiSeq 2500 platform. The gRNA library sequences were used to align the sequencing reads and the counts for each gRNA were determined with MAGeCK. Screens were analyzed with MAGeCK RRA (0.5.9.5), and the two biological screen replicates were analyzed independently (Sup. Table 1). To calculate a single value for the significance of effect, a ratio was taken between the positive and negative MAGeCK scores and log-transformed. The top 36 hits associated with a PDX1 decrease had a MAGeCK neg.FDR<0.1 in replicate 2 and caused a PDX1 decrease in both replicates (Sup. Table 1). Hits overlapped with annotated promoter regions were designated as “Promoter Hits,” with the rest designated as “Non-Promoter Hits.” Promoter overlap was defined as overlap with an interrogated region ± 1.5kb of Gencode (v18) TSS sites. For Figure 1F, if there were multiple overlapped regions, the most significant hit was used for plotting. To link enhancers to the putatively regulated genes, the nearest linearly adjacent “Promoter Hit” was inferred to be the regulated gene of the “Non-Promoter Hit.” To determine hit and non-hit overlap with H3K27ac ChIP-seq peaks during differentiation, Genrich (0.6.1)^103^ was used to call peaks at an FDR-adjusted p-value of 0.01 on previously published ES, DE, GT, and PFG stage H3K27ac ChIP-seq data^25^. Conservation scores of regions were determined by calculating the average of per nucleotide conservation across all nucleotides within each region from PhyloP100 per-nucleotide conservation scores^69^.

### Generation of *ONECUT1e-664kb* clonal knockout hPSC lines

H1 iCas9 hPSC lines^62^ were used to generate three enhancer deletion lines carrying *ONECUT1e-664kb* deletions. Two gRNAs targeting the flanking regions of the enhancer were designed, and knockouts generated as previously described but with some modifications^8^. gRNAs and tracer RNA were ordered from IDT (Alt-R CRISPR-Cas9 crRNA, 1072532) and added at a 15 nM final concentration. In brief, gRNA/tracer RNA and Lipofectamine RNAiMAX (Thermo Fisher Scientific, 13778030) were diluted separately in Opti-MEM (Invitrogen, 31985070), mixed together, and incubated for 15 min at room temperature, and then added dropwise to freshly seeded iCas9 hPSCs in a 24-well plate. DOX (2 μg/ml) was added the day before transfection, the day of transfection, and 1 day after transfection to induce Cas9 expression. Three days after transfection, hPSCs were dissociated into single cells and ∼1**-**3E3 cells were plated into a 10 cm tissue culture dish for colony formation. After ∼10 days of expansion, single colonies were picked. Genomic DNA from crude cell lysate was used for PCR genotyping (primers: ONECUT1e_KOgenot_F, ONECUT1e_KOgenot_R). Heterozygous lines contained a band corresponding to the WT band and a smaller band corresponding the knockout band, while homozygous lines only had smaller bands. Further genotyping was done through TOPO cloning of the genotyping amplicons which determined the exact genotypes of both alleles for all lines (Sup. Table 2). gRNA target sequences and primers used for PCR and sequencing are listed in Supplementary Table 3.

### Generation of clonal hPSC lines with disease-associated variant

A process similar to enhancer deletion line generation was used to generate heterozygous knock-in SNP lines carrying the rs528350911 variant. During transfection, a single gRNA proximal to the variant site was used, and a ssDNA HDR template containing the disease-associated variant was added to the mixture at the Lipofectamine transfection step (Sup. Table 3) at a concentration of 20 nM. Genomic DNA from crude cell lysate was used for PCR (primers: ONECUT1e_KOgenot_F, ONECUT1e_KOgenot_R). All picked clones were sequenced with both primers, and heterozygous lines picked by the presence of a double peak on the spectrogram that corresponded to the insertion of the disease-associated variant. For lines containing the introduced variant, no indels or other changes from the reference genome were detected in the 584bp amplicon.

### *ONECUT1* locus SNP phasing

To phase all SNPs in the *ONECUT1* locus, Nanopore sequencing with adaptive sampling of the region chr15:52,707,645-53,586,740 (and controls regions) was conducted on an H1 iCas9 line with the rs528350911 heterozygous variant (line N11). DNA was extracted using Monarch® HMW DNA Extraction Kit for Cells & Blood (NEB, T3050), according to the manufacturer’s instructions. DNA quantity was measured using a QuantiT (Thermo Fisher Scientific), purity on a NanoDrop (Thermo Fisher Scientific) and fragment-size distribution on a TapeStation (Agilent). Prior to ONT library preparation, DNA was sheared to ∼15-20 kb fragment size using Covaris g-TUBE. Sequencing libraries were prepared from ∼1–2ug of DNA using the native library preparation kit SQK-LSK114 according to the manufacturer’s instructions. Each library was loaded onto a PromethION flow cell R10.4.1 and sequenced on an ONT PromethION P24 device. The sample was run for 72 h with 1 nuclease flush and reload performed during the run to maximize sequencing yield.

Reads were aligned with minimap2 (2.26)^106^, with samtools^107^ coverage estimating ∼138X coverage across the region enriched with adaptive sampling. WhatsHap (2.2)^108^ was used to phase the variants in the locus as well as assemble a haplotype containing the *ONECUT1* gene, enhancer, and surrounding regions (Sup. Fig. 2D). WhatsHap was also used to phase-tag reads for visualization in Sup. Fig. D and E. From these results, a heterozygous SNP was identified in the 3’UTR of the *ONECUT1* transcript, as well as the enhancer-proximal Genotyping SNP. *ONECUT1e-664kb* heterozygous knockout line phasing To determine which allele contained the knocked-out enhancer in the heterozygous enhancer deletion lines (Het. 1, Het. 2), a forward primer was designed centromeric to the Genotyping SNP and a reverse primer targeting the enhancer sequence that would be present on the non-deletion allele. Since line designated as Het. 1 contained an inversion of the WT enhancer sequence, two different reverse primers were needed. Thus, for Het. 1, and amplicon was generated with the primer pair ONECUT1e_GenotSNP_F, ONECUT1e_GenotSNP_R_WT_inv_e, and for Het.2, the pair ONECUT1e_GenotSNP_F, ONECUT1e_GenotSNP_R_WT_e. The amplicon was sent for sanger sequencing with the ONECUT1e_GenotSNP_F primer used as the sequencing primer. The Genotyping SNP from this sequencing was used to determine the allele containing the deletion.

### *ONECUT1e-664kb* heterozygous rs528350911 knock-in line phasing

A PCR amplifying an ∼8.1kb fragment containing both the T2D-associated variant and the Genotyping SNP (primers: ONECUT1e_GenotSNP_F, ONECUT1e_GenotSNP_R_WT_e) with the LongAmp Taq polymerase kit was conducted. The amplicon was sent to Plasmidsaurus for linear fragment sequencing. Reads were aligned with minimap2 (2.26), and variants were phased with longshot (0.4.5)^109^.

### Immunofluorescence staining

PFG-stage pancreatic differentiated cells were fixed in 4% paraformaldehyde (Thermo Fisher Scientific, 50980495) for 10 min at room temperature. After washing with PBST (PBS with 0.1% Triton X-100) three times, cells were blocked in 5% donkey serum in PBST buffer for 30 min at room temperature. Primary and second antibodies were diluted in the blocking solution (Sup. Table 4). Cells were incubated with primary antibodies overnight at 4°C, followed by 1 hr staining for secondary antibodies at room temperature. The cells were then stained with 4′,6-diamidino-2-phenylindole (DAPI) for ∼15 min at room temperature. Images were taken using a confocal laser scanning platform (Leica TCS SP5).

### Flow cytometry analysis

Cells were dissociated using TrypLE Select and resuspended in FACS buffer (5% FBS in PBS). LIVE/DEAD Fixable Violet Dead cell stain (Invitrogen, L34955) was used to discriminate dead cells from live cells. LIVE/DEAD staining was performed per manufacturer’s instructions in FACS buffer, followed by fixation and intracellular staining with a FOXP3 staining buffer set (eBioscience, 00-5523-00) following the manufacturer’s instructions. Permeabilization/fixation was performed at room temperature for 30 minutes. Antibody staining was performed in permeabilization buffer for 30 minutes. Antibodies for this study are listed in Supplementary Table 8. Cells were then analyzed using BD LSRFortessa. Flow cytometry analysis and figures were generated using FlowJo v.10.

### RNA-seq and analysis

RNA was extracted from differentiated cells with the Zymo Quick-RNA MiniPrep kit (R1055). TruSeq stranded total (Ribodepletion) RNA-seq was performed on three independent differentiations by the MSKCC Integrated Genomics Operation (IGO) Core. Briefly, STAR alignment was performed followed by featureCounts^110^ quantification (Sup. Table 6) and normalization with DESeq2 median of ratios prior to visualization in Fig. 3A, Fig. 3C, and Sup. Fig. 3B. Differential gene expression was performed and adjusted p-values computed with DESeq2. Strand-specific analysis of promoter antisense transcription analysis was performed with featureCounts strand specific quantification of a region telomeric to *ONECUT1* (chr15:52805588-53391232). For strand-specific transcript visualization, samtools was used to extract sense and antisense alignments from .bam files from which bigwig coverage files were generated.

### Droplet digital PCR (ddPCR) and analysis

Assays specific for the detection of SNPs (Sup. Table 5) were designed (using Primer3Plus, if needed) and ordered through Bio-Rad. Cycling conditions were tested to ensure optimal annealing/extension temperature as well as optimal separation of positive from empty droplets. Optimization was done with a known positive control. A constant amount of RNA or DNA was used within each experiment. Reactions were partitioned into a median of ∼17,500 droplets per well using the QX200 droplet generator. Plates were read and files analyzed with the QuantaSoft software to assess the number of positive droplets.

Procedure for RNA: after PicoGreen quantification of RNA, droplet generation was performed on a QX200 ddPCR system (Bio-Rad catalog # 1864001) using cDNA generated from 0.1-2 ng total RNA with the One-Step RT-ddPCR Advanced Kit for Probes (Bio-Rad catalog # 1864021) according to the manufacturer’s protocol with reverse transcription at 42°C and annealing/extension at 60°C.

Procedure for DNA: after PicoGreen quantification of DNA, 0.25-9 ng gDNA of ChIP-ed DNA was combined with locus-specific primers; FAM-and HEX-labeled probes; Hae III (DNA extracted form hPSC SNP lines), Hind III (ChIP-ed DNA), or Mse I (DNA extracted *ONECUT1e-664kb* hPSC knockout lines); and digital PCR Supermix for probes (no dUTP). Emulsified PCRs were run on a 96-well thermal cycler using cycling conditions identified during the optimization step (95°C 10’; 40 cycles of 94°C 30’ and 53°C (rs528350911 assays only) or 60°C 1’; 98°C 10’; 4°C hold).

### Hi-C and analysis

Hi-C was performed as previously described with cells differentiated to the PFG stage processed with the Arima Hi-C kit (Arima, A510008)^8^. HiC-Pro (3.1.0)^111^ was used to align the individual Hi-C replicates to GRCh38, GCA_000001405.15, with alternative contigs removed, and then final merged libraries were obtained by combining all *.allValidPairs files where duplicates across biological replicates are maintained. DE libraries consisted of three biological replicates (two HUES8, one H1 with accessions 4DNFIKCEWMIA.hic, 4DNFI7RGXYFY.hic, and 4DNFIRLG5UWL.hic respectively), and the PFG libraries consisted of two biological replicates differentiated to the PFG stage (merged accession 4DNFIQBPRZZD.hic). For visualization, Hi-C data was visualized at 25 KB resolution.

Virtual 4C counts for the regions of interest were extracted using utilities from plotGardener (1.6.4)^112^ and plotted as a one-dimensional signal. Hi-C matrix counts for the regions of interest were extracted using plotGardener and plotted as a rectangular region. Observed counts are library normalized prior to plotting so that the total number of reads between panels is comparable. TADs that were previously identified were also overlaid on the data and can be found in the Hi-C subseries of GEO data set published with this work. Counts within the vicinity of the *ONECUT1* promoter (all contacts whose start or end bins sit within a 1.2Mb window around the *ONECUT1* promoter), were extracted with strawr (0.0.91) and passed to DESeq2 (1.40.1)^113^. The Wald test was used to assess significance of all differential contacts within that window, and significant hits anchored on the *ONECUT1* containing bin are reported at an FDR of 0.1.

For calling topologically associating domains, merged Hi-C files were first ICE normalized through HiTC (1.34.0)^114^ and then topologically associating domains were identified with HiCDCplus (1.5.2)^115^ using its interface to TopDom at 50 kb. Window size for scanning was set at 5.

### ChIP and ChIP-seq assays

ChIP was performed as previously described^18^. For each sample, around 30E6 cells were crosslinked in-plate with 1% formaldehyde for 10 minutes at 37°C and quenched with 0.125 M glycine. Fixed cells were collected and washed in cold PBS buffer, snap frozen, and saved at - 80°C. On the day of immunoprecipitation, the cell pellet was thawed on ice, resuspended in 700uL SDS buffer (1% SDS, 10 mM EDTA, 50 mM Tris–HCl, pH 8), and incubated for 10min on ice. Sonication was performed on a Branson Sonifier 150 set at 30% amplitude for 5.5min (10s on/off pulsing). Supernatant was pre-cleared with Dynabeads Protein G (#10004D) and then incubated overnight with antibodies (10 ug for transcription factors, 5 ug for H3K27ac, Sup. Table 4) at 4°C.). On the next day, the ChIP samples were incubated with Dynabeads Protein G for 6 hours at 4°C. Then the beads were pelleted and sequentially washed twice with low salt (0.1% SDS, 1% Triton X-100, 2 mM EDTA, 20 mM Tris–HCl, pH 8, 150 mM NaCl), then high salt (0.1% SDS, 1% Triton X-100, 2 mM EDTA, 20 mM Tris–HCl, pH 8, 500 mM NaCl), and finally TE buffer (10 mM Tris–HCl, pH 8, 1 mM EDTA). The DNA was eluted from the beads by incubating in elution buffer (1% SDS, 0.1 M NaHCO3) at 65°C for 15min and decrosslinked with 190 mM NaCl at 65°C overnight. A total of 5 μl 0.5 M EDTA, 10 μl 1 M Tris–HCl, pH 6.5, and 1 μl Proteinase K (20 mg/ml) were added to 260 uL of the de-crosslinked product and incubated for 1 hr at 45°C. DNA was isolated by using QIAquick PCR purification kit (Qiagen, 28104; QIAGEN). H3K27ac ChIP samples were submitted to MSKCC Integrated Genomics Operation core for NGS library preparation and sequencing. GATA4, GATA6, and FOXA2 ChIP samples were submitted for ddPCR.

### H3K27ac ChIP-seq Analysis

Sequenced data were aligned to the hg38 reference genome using bowtie2 (2.5.1)^116^. Peak calling was performed using MACS2 (2.2.9.1)^117^ with the respective ChIP input as the control and the default P value cutoff and default extension size. Irreproducible discovery rate (IDR)^118^ with a cutoff of 0.01 was used to filter reproducible peaks, where at least 2 pairwise comparisons of the three replicates passed the cutoff. Peaks were filtered by ENCODE exclusion list version 2 for hg38^119^. The quantification and differential signal intensity of the samples were calculated in DESeq2^113^. The integer counts, log(fold-change), and adjusted P values are provided in Supplementary Table 7. The replicates were combined, and the signal track was generated from MACS2. MACS2 subtract was used to generate the difference of the signal track. All signal tracks were visualized using Integrative Genomics Viewer (IGV)^120^.

For allele-specific analysis, GATK^121^ ASEReadCounter was used to quantify reads at heterozygous positions phased by long-read sequencing and ratios were calculated for heterozygous positions (score filters: PHASE_QUAL >= 140, and QUAL >= 200) with more than 100 reads total across all WT and Het. 2 replicates. For Figure 4F, the “Phased Region” is the haplotype block containing *ONECUT1* called by WhatsHap (chr15:50462878-54550826) and visualized in Sup. Fig. 2I.

### chromBPNet variant modeling

A ChromBPNet^81,82^ model was trained to predict the local ATAC profile using bulk ATAC-seq data from the PFG stage^25^. Default hyperparameters were utilized with an input size of 2114 base pairs and an output size of 1000 base pairs for training both the bias model and the ChromBPNet model. For the bias model training, regions were selected that do not overlap with any ATAC-seq peaks in the four stages of pancreas differentiation^25^, applying a bias threshold factor of 0.3 to filter out high-count regions in the sets. This resulted in a training set of 54957 regions. For ChromBPNet training, ATAC-seq peaks only from the PFG stage with IDR values greater than 830 were used, resulting in a training set of 52541 regions. During training for both models, peaks from chromosomes 15 and 13 for validation were held out. For each specific region of interest, the 2114 base pairs around the summit center was used as input and the DeepLift attribution score obtained using the trained ChromBPNet model. Specifically, the original attribution pipeline from the ChromBPNet GitHub repository was employed, which involves computing the DeepLift score over 20 reference inputs shuffled from the original sequence. To find the GATA motifs, FIMO^122^ was used on our region of interest using the positional weight matrix (PWMs) of all GATA variants from the CIS-BP2.00 dataset^123^, and motif hits with p-values < 0.05 were filtered out.

## QUANTIFICATION AND STATISTICAL ANALYSIS

All datapoints refer to biological repeats. No statistical method was used to predetermine sample sizes. The investigators were not blinded to allocation during experiments and outcome assessment. No data were excluded from the analyses unless the differentiation experiment itself failed. The number of biological and technical replicates are reported in the legend of each figure. Flow cytometry analysis was derived from at least three independent experiments. For ChIP-seq and bulk RNA-seq, quantification and statistics were derived from at least three independent experiments. CRISPR dCas9-KRAB screening was performed twice. Quantification is shown as the mean ± s.d. All the statistical analysis methods are indicated in the figure legends and methods.

## Supplemental Information

### Supplementary Figures

**Sup. Fig. 1, Related to Figure 1:**
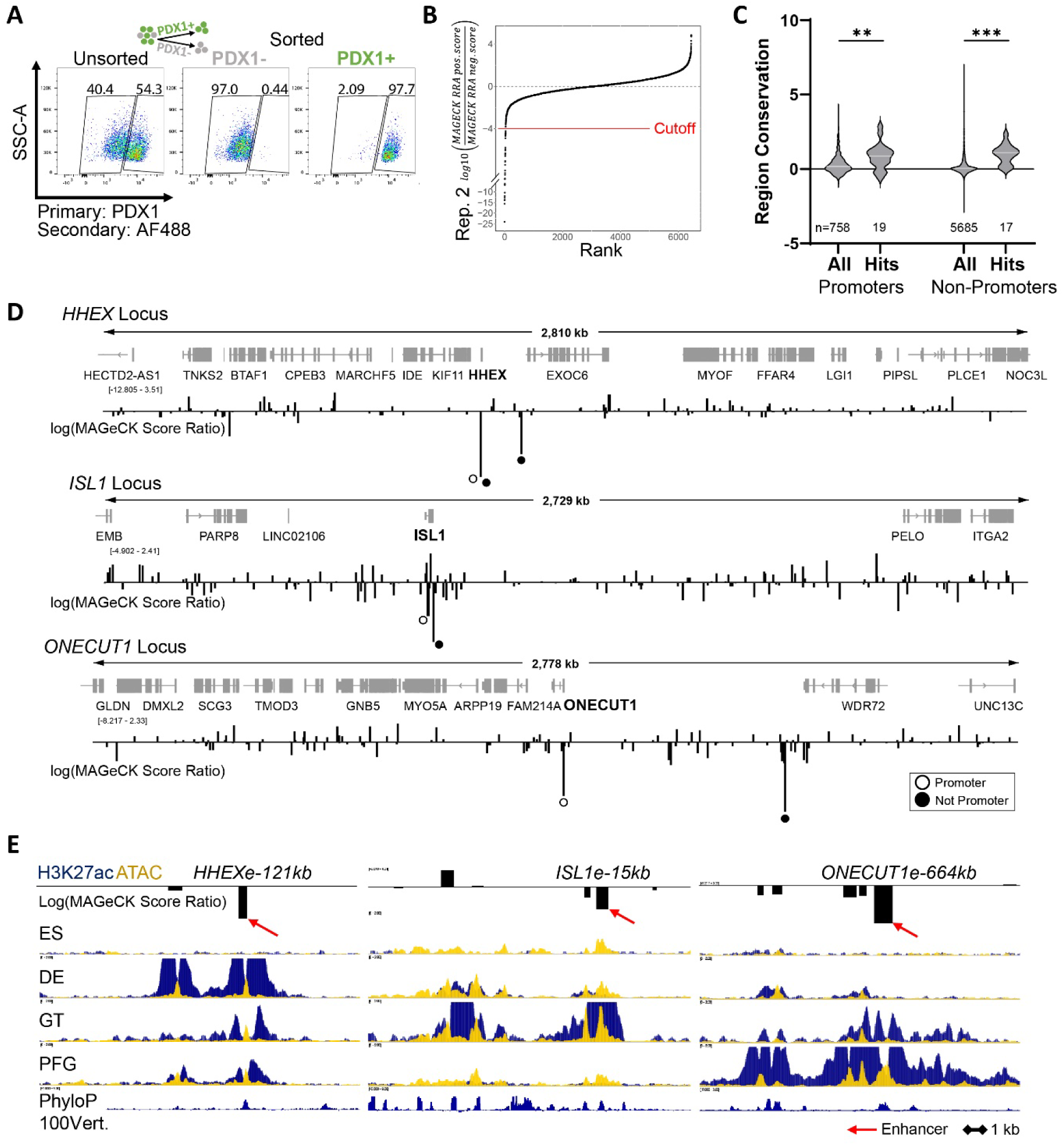
Supplemental results and analysis for CRISPRi screens. A) Representative flow cytometry plot of PFG differentiated cells pre- and post-sort PDX1 sorting. B) Rank plot of replicate 2 MAGeCK scores. C) Calculated conservation scores for each region within indicated group. Ordinary one-way ANOVA with Šidák’s multiple comparisons test, with a single pooled variance. D) Representative IGV plots for enhancer-gene pair assignment rationale. Each bar represents a MAGeCK score ratio from replicate 2. A negative bar indicates region repression caused a decrease in the proportion of PDX1+ cells, with bar size indicating the magnitude of the change in PDX1 enrichment. Hollow black points represent promoter region hits, filled black points represent non-promoter region hits. E) H3K27ac ChIP-seq, ATAC-seq, and phyloP100way vertebrate conservation at selected enhancer loci.

**Sup. Fig. 2, Related to Figure 2:**
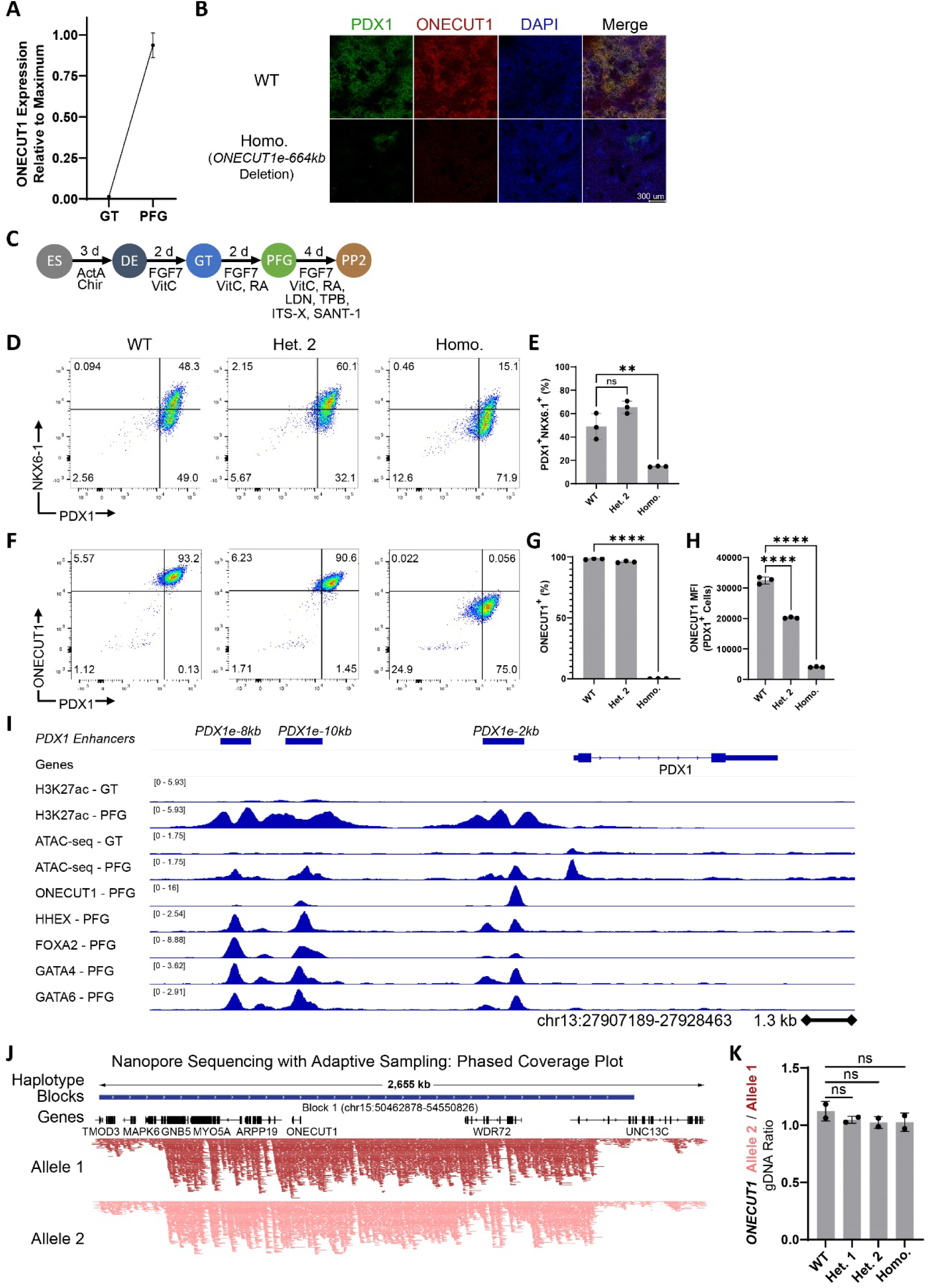
*ONECUT1e-664kb* deletion supplemental results and analysis. A) ONECUT1 transcript ddPCR on RNA from WT cells at GT and PFG stages. B) PFG-stage immunofluorescence for PDX1 and ONECUT1 of WT and homozygous enhancer knockout cells. C) ES to PP2 hPSC stepwise pancreatic differentiation protocol schematic. D) Representative flow cytometry plots of WT and enhancer knockout cells differentiated to the PP2 stage. E) Percentage of cells achieving a PDX1+NKX6.1+ identity. For panels E, G, and H, each dot represents one independent experiment (n = 3 independent experiments) and data are presented as the mean ± s.d. One-way analysis of variance (ANOVA) followed by Dunnett multiple comparisons test versus WT control. F) Representative flow cytometry plots of WT and enhancer knockout cells differentiated to the PP2 stage. G) Percentage of cells achieving a ONECUT1+ identity. H) ONECUT1 MFI quantification from flow cytometry of unedited and enhancer knockout cells differentiated to the PP2 stage. I) Transcription factor binding and epigenetic profile of the PDX1 locus in pancreatic differentiation. ONECUT1 and HHEX ChIP-seq from (Yang et al., 2022, GSE181480) and FOXA2, GATA4, and GATA6 ChIP-seq from (Lee et al., 2019, GSE114102). J) Plot of phased reads and haplotype blocks from nanopore sequencing of H1 hPSCs with adaptive sampling. K) ONECUT1 transcript ddPCR on DNA from WT and enhancer knockout cells. One-way analysis of variance (ANOVA) followed by Dunnett multiple comparisons test versus WT control.

**Sup. Fig. 3, Related to Figure 3:**
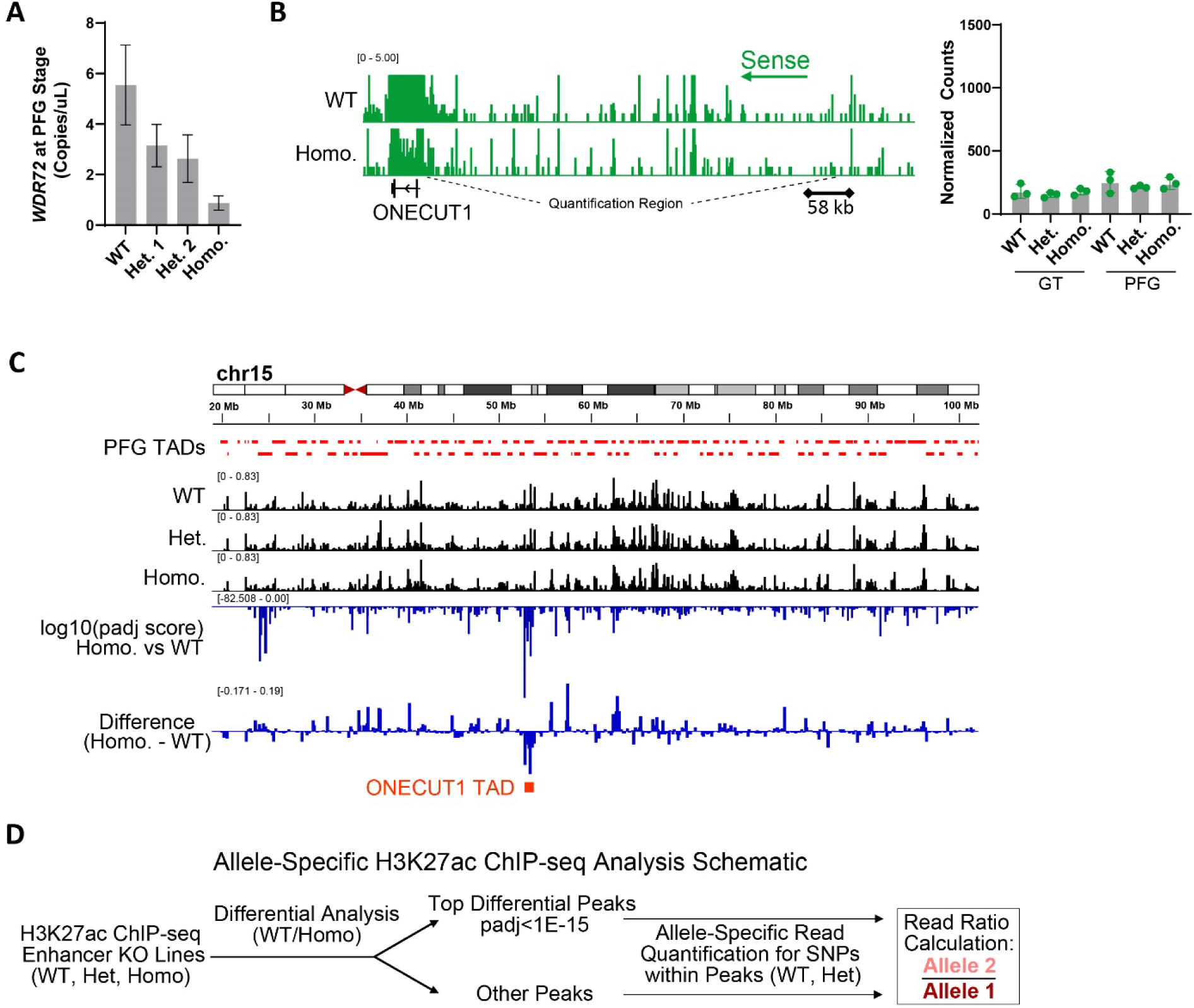
Supplemental RNA-seq and H3K27ac ChIP-seq results and analysis. A) Non allele-specific WDR72 ddPCR results of WT and enhancer knockout cells differentiated to the PFG stage. B) Visualization of sense read coverage (relative to ONECUT1) in region telomeric to ONECUT1 from strand-specific RNA-seq of PFG-stage WT and homozygous enhancer knockout cells (representative differentiation replicate). Each dot represents one independent experiment (n = 3 independent experiments) and data are presented as the mean ± s.d. One-way analysis of variance (ANOVA) followed by Dunnett multiple comparisons test versus WT control. C) Full chromosome view of H3K27ac in WT and enhancer knockout cells showing the ONECUT1 TAD to have the most significantly affected regions, and the regions with the greatest magnitude loss of H3K27ac. D) Allele-specific H3K27ac ChIP-seq analysis schematic.

**Sup. Fig. 4, Related to Figure 4:**
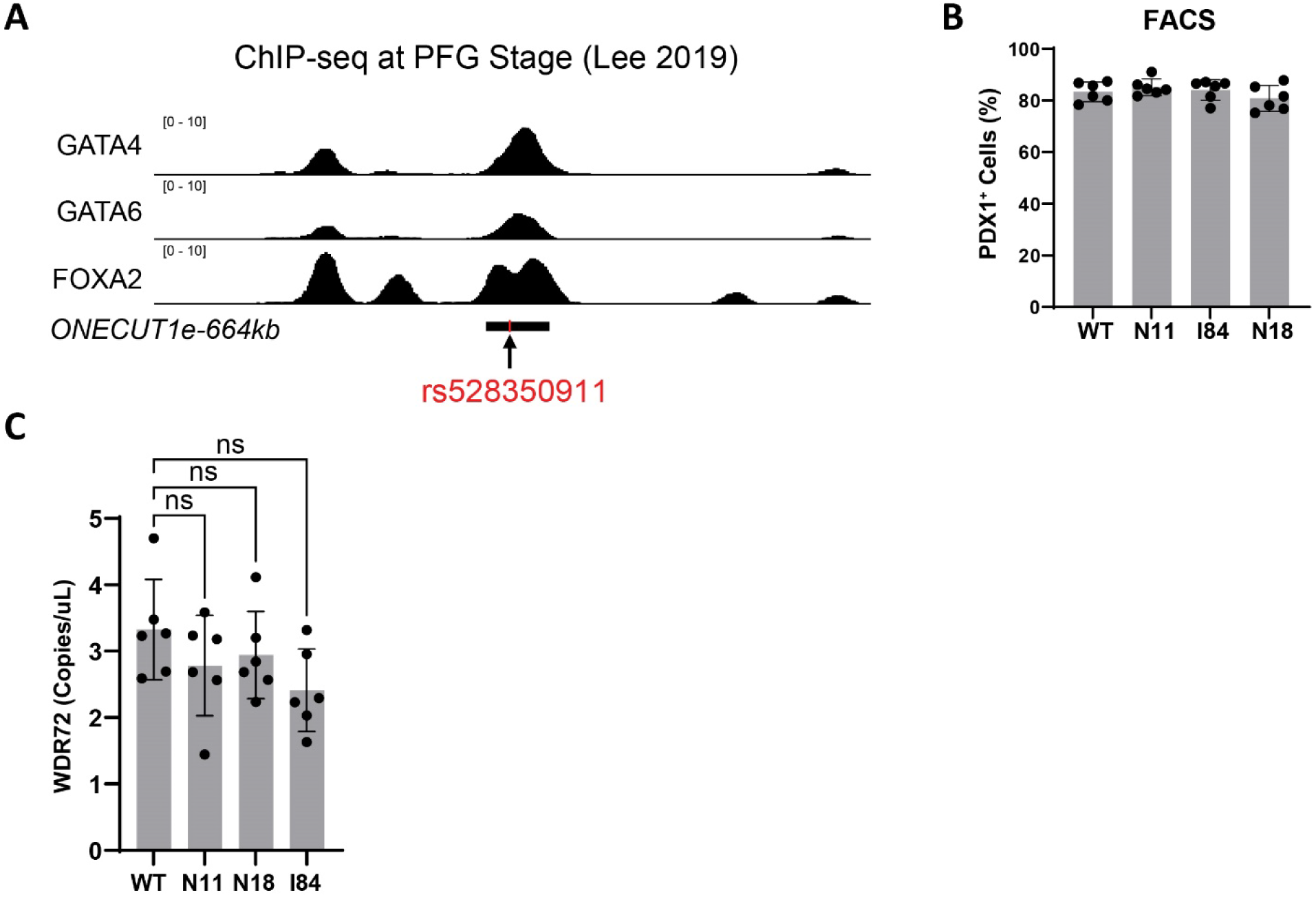
Supplemental results for T2D-associated SNP hPSC modeling. A) Visualization of ChIP-seq at ONECUT1e-664kb locus conducted at PFG stage. Black bar indicates ONECUT1e-664kb. B) Bar plots of percentage of cells achieving a PDX1+ identity. Each symbol represents one independent experiment (n = 6 independent experiments) and data are presented as the mean ± s.d. One-way analysis of variance (ANOVA) followed by Dunnett multiple comparisons test versus WT control. C) Non allele-specific WDR72 ddPCR results of WT and enhancer SNP cell lines differentiated to the PFG stage.

## Supplementary Tables

**Table S1:** Enhancer screen region selection and analysis results. Sheet 1: Transcription factor loci selected for CRISPRi interrogation. Sheet 2: ATAC-seq accessible regions selected for interrogation. Sheet 3: Read abundance quantification of each gRNA. Sheet 4: MAGeCK-RRA region-level analysis of two screens. Sheet 5: Top screen hits, FDR<0.1 in screen 2.

**Table S2:** Engineered hPSC line allele genotypes.

**Table S3:** List of gRNA target sequences and primer sequences for PCR genotyping.

**Table S4:** Antibody target, company or lab that supplied antibody, catalog number, application, and quantity or dilution used are indicated. For ChIP experiments, the amount used for cells from around half of a 15cm plate (roughly 40-50 million cells) is indicated.

**Table S5:** ddPCR assay details.

**Table S6:** RNA-seq conducted on WT, heterozygous (het. 2) and homozygous *ONECUT1e-664kb* enhancer knockout cells. Sheet 1: DESeq2 normalized gene counts. Sheet 2: DESeq2 differential analysis between WT and heterozygous enhancer knockout cells at PFG stage. Sheet 3: DESeq2 differential analysis between WT and homozygous enhancer knockout cells at PFG stage.

**Table S7:** H3K27ac ChIP-seq conducted on PFG-stage WT, heterozygous (het. 2), and homozygous *ONECUT1e-664kb* enhancer knockout cells. Sheet 1: Raw integer counts of H3K27ac reads. Sheet 2: DESeq2 calculated differential statistics between pairwise comparisons among WT, heterozygous, and homozygous enhancer knockout cells. Sheet 3: Phased read counts at indicated position, read ratios for each replicate, average read ratios for each genotype. Top_Differential_Peak indicates whether the position falls within a top differential peak (WT vs Homo, padj>1E-15).

